# Dynamic information sharing across frontal-basal ganglia neurons during cost-benefit decision-making and reward pursuit

**DOI:** 10.64898/2026.11.13.766393

**Authors:** Oliver Härmson, Isaac Grennan, Brook Perry, Robert Toth, Mark E. Walton, Sanjay G. Manohar, Andrew Sharott

## Abstract

Cost–benefit decision-making requires animals to integrate prospective rewards with the costs of obtaining them and translate these evaluations into motivated action. How population activity is coordinated across the frontal–basal ganglia network during these processes remains incompletely understood. Here, we recorded simultaneously from the anterior cingulate cortex, medial/ventral orbitofrontal cortex, dorsomedial striatum, ventral pallidum and subthalamic area in rats performing a cost–benefit decision task. Using canonical correlation analysis, we identified shared population activity across these anatomically distributed circuits and characterised its dimensionality, task-related information and relationship to behavioural sensitivity. The strength and dimensionality of inter-area coupling varied across task epochs and circuit classes, with frontal–basal ganglia interactions exhibiting particularly high dimensionality during reward pursuit. Shared activity represented reward and time on task (trial count) during decision-making, while movement-related information became increasingly prominent during reward pursuit. These representations also depended on subsequent choice: reward was more strongly represented on accepted trials, whereas time on task was more prominent on rejected trials. The strength of inter-area coupling was further associated with behavioural sensitivity across sessions. Stronger coupling between the subthalamic area and the dorsomedial striatum was associated with reduced reward sensitivity. By contrast, communication between the medial/ventral orbitofrontal cortex and dorsomedial striatal was associated with reduced effort aversion. Together, these findings reveal a dynamic, circuit-specific organisation of shared neural activity during cost–benefit behaviour and link inter-area coupling to variation in motivated choice. They suggest that distinct communication pathways may contribute to specific facets of motivated choice, with applications to targeted neuromodulation therapies for disorders of motivation.

## Introduction

Cost–benefit decision-making depends on distributed neural computations that integrate the expected benefits of an action with the costs required to obtain them. Across frontal cortical and basal ganglia regions, single-neuron firing rates have been shown to encode variables including reward magnitude, effort cost, action value and movement-related signals (Härmson et al., 2025; Hunt et al., 2018; Kennerley & Wallis, 2009; Nougaret et al., 2022; Tachibana & Hikosaka, 2012). Such representations are spatially distributed, with related decision variables often expressed simultaneously across multiple interconnected structures. At the population level, ensemble activity within, and in some cases spanning, individual brain regions can likewise contain information about the variables that guide choice and subsequent behaviour (Härmson et al., 2025). Together, these findings suggest that cost–benefit decisions may emerge from coordinated activity across a broader frontal–basal ganglia network rather than from computations confined to any single region.

How this distributed activity is coordinated across regions, however, remains less well understood. Anatomical and disconnection studies indicate that interactions between the frontal cortex and basal ganglia are necessary for appropriate value-guided behaviour (Christakou et al., 2004; Hart et al., 2018; Hauber & Sommer, 2009; Jenni et al., 2022; Oyama et al., 2022). Large-scale single neuron recordings in primates have shown dynamic inter-regional communication between areas of the frontal cortex during value-guided behaviour (Stoll & Rudebeck, 2024). Yet the extent to which shared activity across frontal–basal ganglia structures underpins cost–benefit decision-making has received comparatively little attention. For instance, it is unknown whether information is shared via one or multiple latent dimensions and whether these are used to transmit internal or external cost-benefit decision-relevant information. Of clinical relevance to syndromes such as apathy and anhedonia, it is not known whether such shared activity can predict moment-to-moment decisions or longer-term behavioural sensitivity to reward or effort. Characterising the information carried by shared cross-area activity may therefore advance our understanding of the circuit-level organisation of motivated behaviour and help identify candidate pathways for targeted neuromodulation in disorders of motivation.

Here, we investigated cross-area population coupling within a set of anatomically distributed regions important for cost-benefit decision-making, comprised of the anterior cingulate cortex (ACC), medial/ventral orbitofrontal cortex (MO/VO), dorsomedial striatum (DMS), ventral pallidum (VP) and subthalamic area (STA) while rats performed a previously established cost–benefit decision task (Härmson et al., 2025). By applying canonical correlation analysis, we found that cost-benefit decision-making and reward pursuit was accompanied by dynamic changes in the strength and dimensionality of shared population activity across frontal–basal ganglia and intrinsic basal ganglia circuits. These shared representations carried temporally and anatomically specific information about reward, internal state and movement, with reward-related coding differing according to whether offers were accepted or rejected. Moreover, the strength of coupling in distinct circuits was associated with reward and effort sensitivity within animals. Together, these findings identify dynamic, circuit-specific patterns of shared neural activity that are associated both with the information available during cost–benefit decisions and with individual variation in motivated behaviour.

## Results

### Cost-benefit decision-making and reward pursuit is paralleled by dynamic communication changes in frontal-basal ganglia circuits

To investigate how distributed neural populations interact during cost–benefit decision-making and reward pursuit, we recorded activity simultaneously from a distributed set of frontal cortical and basal ganglia regions while rats performed a cost-benefit task (fig. 1a-b, S1a-c). In this task rats decided whether to traverse a corridor to a reward magazine as a function of the offered reward size and the travel cost. Reward offers were varied pseudo-randomly on a trial-by-trial basis while effort varied in blocks of ∼20 trials (STAR methods). Analyses focused on accepted trials (i.e. trials in which animals collected the offered reward) in discrete task epochs spanning task initiation (pre-cue), offer presentation (offer), action initiation (action), reward pursuit (corridor light-gate 1 and 6 crossing), and the pre-reward delay period. We used canonical correlation analysis (CCA) to identify low-dimensional patterns of population activity that were shared between pairs of recorded brain areas (fig. 1c). For each area pair, CCA identified weighted combinations of neuronal firing whose trial-by-trial activity was maximally correlated. We extracted and cross-validated the first 10 canonical dimensions and used permutation testing to determine which dimensions represented statistically reliable inter-area coupling. This framework allowed us to quantify both the strength and dimensional structure of shared population activity across distinct anatomical circuits and behavioural time windows.

**Fig. 1.**
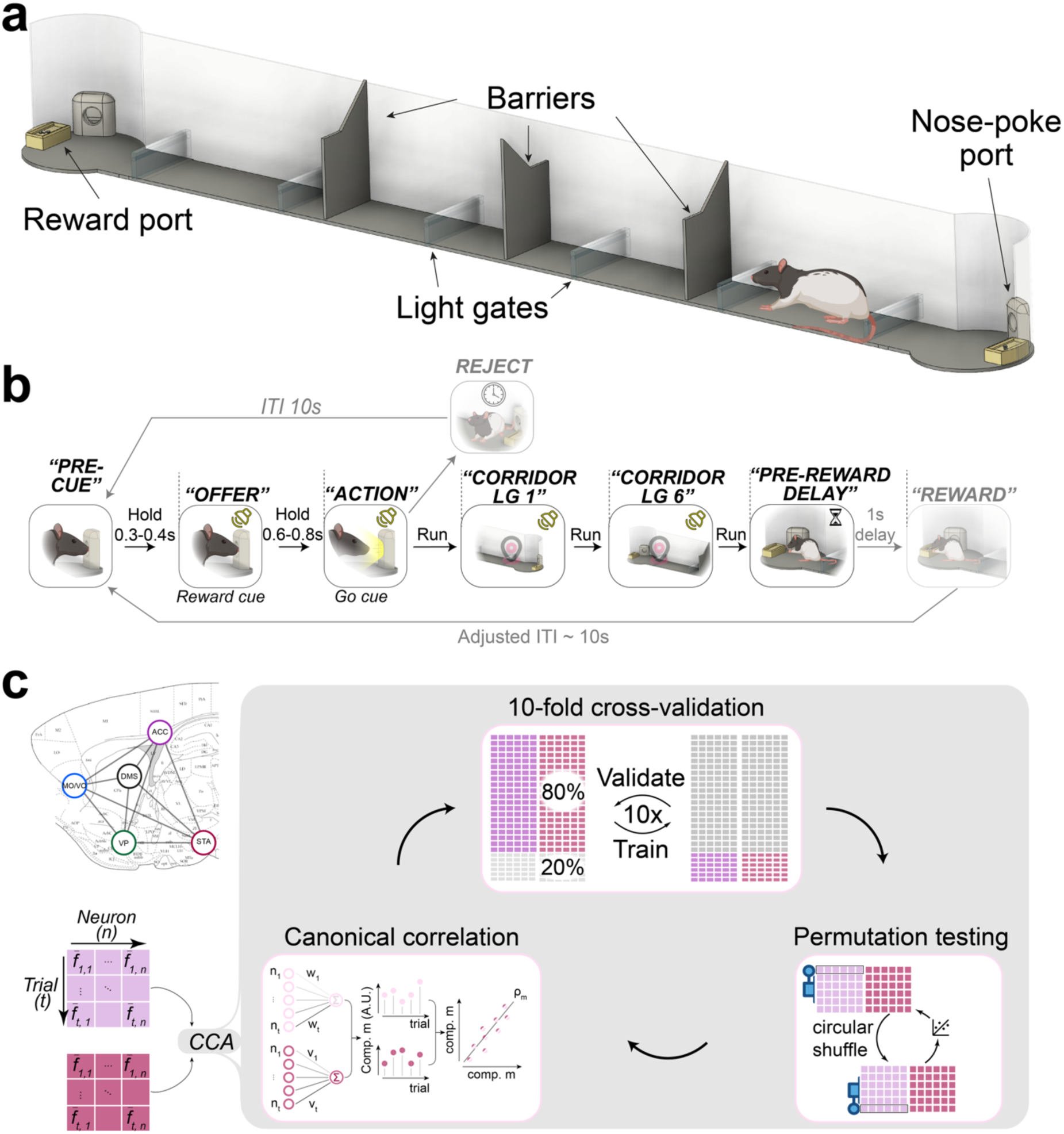
Schematic of cost-benefit task and CCA. **a, Illustration of the experimental apparatus.** This consisted of a 130cm corridor with a variable number of barriers with circular arenas at either end, each of which contained a nose-poke port and a reward port. **b, schematic of the behavioural task and epoch selection.** Rats chose between reward, signalled by an auditory cue during the “offer” window while the rat was in the nose-poke port, available at the far end of a linear corridor (’accept’) versus opting out (’reject’). In this paper, only accepted trials were analysed. Firing data included the first 300ms of the pre-cue period, the first 600ms of the cue period, the first 500ms of the action period, and the first 500ms of the pre-reward delay. Firing rate data during corridor light-gate (LG) 1 and 6 entry included a 500ms centred around the time point of crossing. **c, overview of the CCA analysis pipeline.** Electrophysiology data were histologically verified and spike-sorted as described in (Härmson et al., 2025) and binned into 25ms windows. Firing rates were smoothed with a one-sided exponential decay kernel (τ = 50 ms). The resulting data were from each pair of areas was entered into a CCA pipeline using the python scikit-learn package (STAR methods). The resulting canonical correlation component vectors were trial-by-trial sums of weighted activity from each area that maximised the correlation of aggregate firing rates between the areas. The first 10 canonical component dimensions were extracted where there were sufficient neurons in either area. To avoid overfitting, CCAs were 10-fold cross-validated. Statistical significance was calculated using permutation testing, by shuffling trial-wise activity of area A relative to area B in each CCA model 1,000 times.

We analysed 13,317 trials from 70 sessions in four rats (gh001, gh009, gh010, oh011: 17, 20, 19 and 14 sessions; on average 164 trials per session) (Härmson et al., 2025). This yielded a per-session median and inter-quartile range of 15 (9.5-20.0) units in MO/VO, 8.0 (4.0-13.0) units in ACC, 16.0 (11.5-23.0) units in DMS, 10.0 (9.0-12.0) units in VP, 5.0 (4.0-10.0) units in STA (fig. S1a-c). We first probed whether cross-area coupling varied in relation to the behavioural states on the task. In relation to this, we also asked whether this activity was confined to specific functional sub-circuits in the frontal-basal ganglia network. To address these questions, area pairs were grouped into frontal–frontal (Fr–Fr, comprised here of the MO/VO-ACC pair), frontal–basal ganglia (Fr–BG, encompassing MO/VO or ACC pairings with either the VP, DMS, or STA), and intrinsic basal ganglia (BG–BG, comprised of pairings between DMS, VP and STA) circuit classes. Significant communication channels were identified in each circuit class across the task (fig. 2a). Canonical correlation coefficients in the dominant dimension were consistent between animals across area pairs (fig. S2a–c). The exceptions were MO/VO–ACC and MO/VO–VP, which were recorded in few sessions from few animals (fig. S1c), so their between-animal consistency could not be estimated reliably. Their canonical correlations are nonetheless reported in the main figures. The number of significant CCA dimensions increased during decision-making and reward pursuit, particularly in the frontal-basal ganglia subcircuit (fig. 2a-b). This was supported by a significant circuit group × epoch interaction in a two-way repeated-measures ANOVA (fig. 2b). This was not driven by differences between the combined per-session unit yields between Fr-Fr, Fr-BG and BG-BG pairs (linear mixed model of log unit yield with random intercepts for animal and day; likelihood-ratio test χ²(2) = 1.71, p = .43). Taken together, cost-benefit decision-making and reward pursuit behaviours are accompanied by the emergence of spatially separable communication channels within the frontal-basal ganglia network, with particularly high signal dimensionality in the frontal-basal ganglia circuit.

**Fig. 2.**
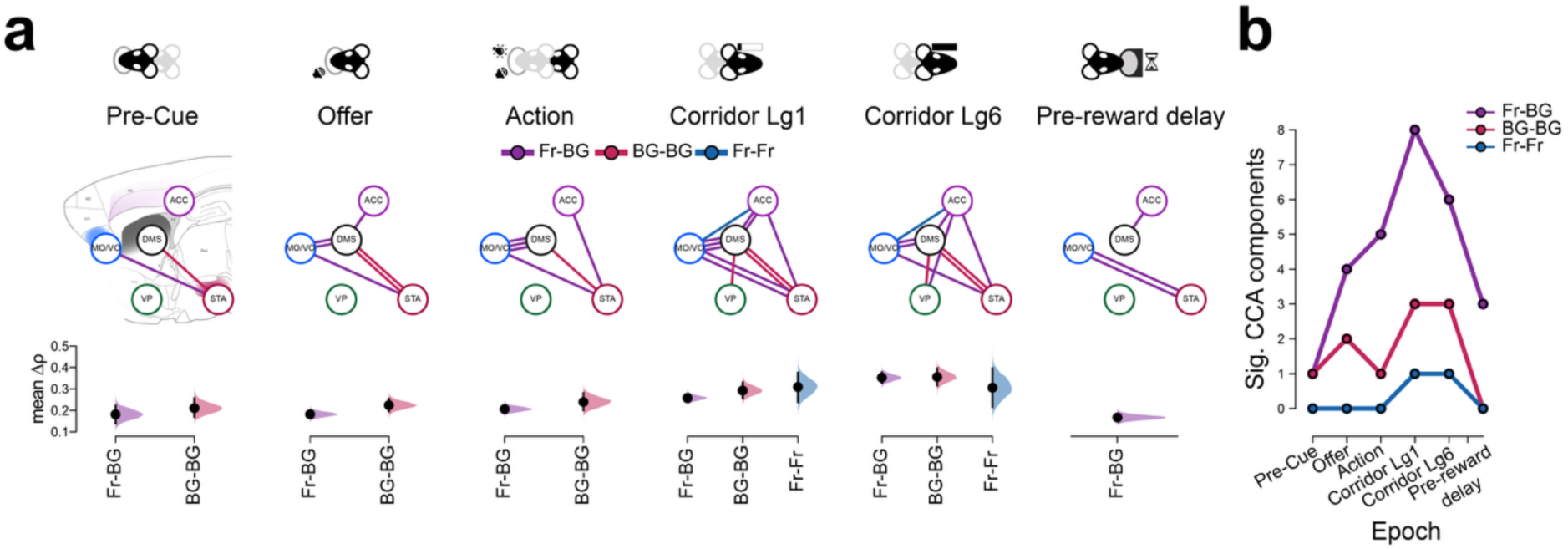
Cortical-basal ganglia areas communicate dynamically through multiple channels during cost-benefit decision-making and reward pursuit. **a,** Node diagrams represent significant inter-area Pearson correlation coefficients (ρ) from the first ten CCA components. Top: nodes represent brain areas, while each edge (line) corresponds to an individual significant CCA dimension. The presence of multiple parallel in an area pair edge indicates that the shared space occupied more than one CCA dimension. Edge widths are fixed. Edges are not represented where the original CCA estimates are not significantly different to their permuted counterparts (one-tailed Wilcoxon test comparing original and mean of permuted datasets across all recorded recording days with a threshold of p < .01). False discovery rate (FDR) was corrected across epochs and area pairs using the Benjamini-Hochberg adjustment (BHA). Bottom: Variance of the ρ estimates in each circuit class per epoch. Δρ values (session by session difference between the original and permuted value) from all significant edges were pooled across sessions and bootstrap-resampled (5,000 resamples). Filled curves show the resulting sampling distribution of the mean, black dots the observed mean, and vertical bars the 95% confidence interval (CI). b, Number of significant canonical-correlation components in each circuit class and task epoch. Coloured lines depict the number of significant dimensions in a particular circuit (colours aligned with panel a) over task epochs (x-axis). The y-axis values are a sum of all significant edges pooled across sessions and are directly inferable from panel a. A 2-way within-subject repeated measures ANOVA with recording session as the repeated measure revealed a significant main effect of circuit type (F_2, 26_ = 26.64, p < .001, η²_G_ = 0.22), epoch (F_5, 65_ = 61.67, p < .001, η²_G_ = 0.33) and a circuit x epoch interaction (F_10, 130_ = 5.28, p = .006, η²_G_ = 0.16). Benjamini-Hochberg-corrected *post-hoc* comparisons indicated that Fr–BG circuits exhibited significantly elevated communication dimensionality within the same circuit during reward-pursuit epochs (corridor lg1 and lg6) relative to pre-cue and pre-reward delay epochs, as well as relative to frontal–frontal (Fr–Fr) dimensionality during reward pursuit epochs (all p<.01; Hedges’ g ≈ 1.42-2.80).

### Shared inter-area activity in the frontal-basal ganglia network represents cost-benefit decision variables in a decision-selective manner

Given the spatiotemporal organisation of communication in the frontal-basal ganglia network, we hypothesised that specific information relating to the task and/or the animal’s internal state is being represented. We thus decoded each of the four key task variables (reward, effort, trial count, or instantaneous velocity) individually from activity in the significant canonical correlation subspace, and compared held-out decoding performance (R^2^) to permuted datasets on a session-by-session basis (STAR methods). There was no significant task-relevant coding present in the pre-cue epoch (fig. 3a-c). This was followed by frequent and strong communication of reward information in the Fr-BG and BG-BG circuits during decision-making and reward pursuit epochs. Reward coding was paralleled and later superseded by trial count and/or velocity coding in those same sub-circuits during decision-making and pursuit epochs, respectively (fig. 3a-c). Where velocity or trial count representations paralleled or replaced earlier reward information within an area pair, these mostly occupied an orthogonal axis (fig. S3a). Of note, reward pursuit epochs were characterised by a higher dynamic range of velocity observations (fig. S3b). By contrast, effort could not reliably decoded from the low-dimensional shared activity between any area pair. Together, this demonstrates the recruitment of specific circuit motifs to represent time-sensitive cost-benefit and movement information during decision making and reward pursuit.

**Fig. 3.**
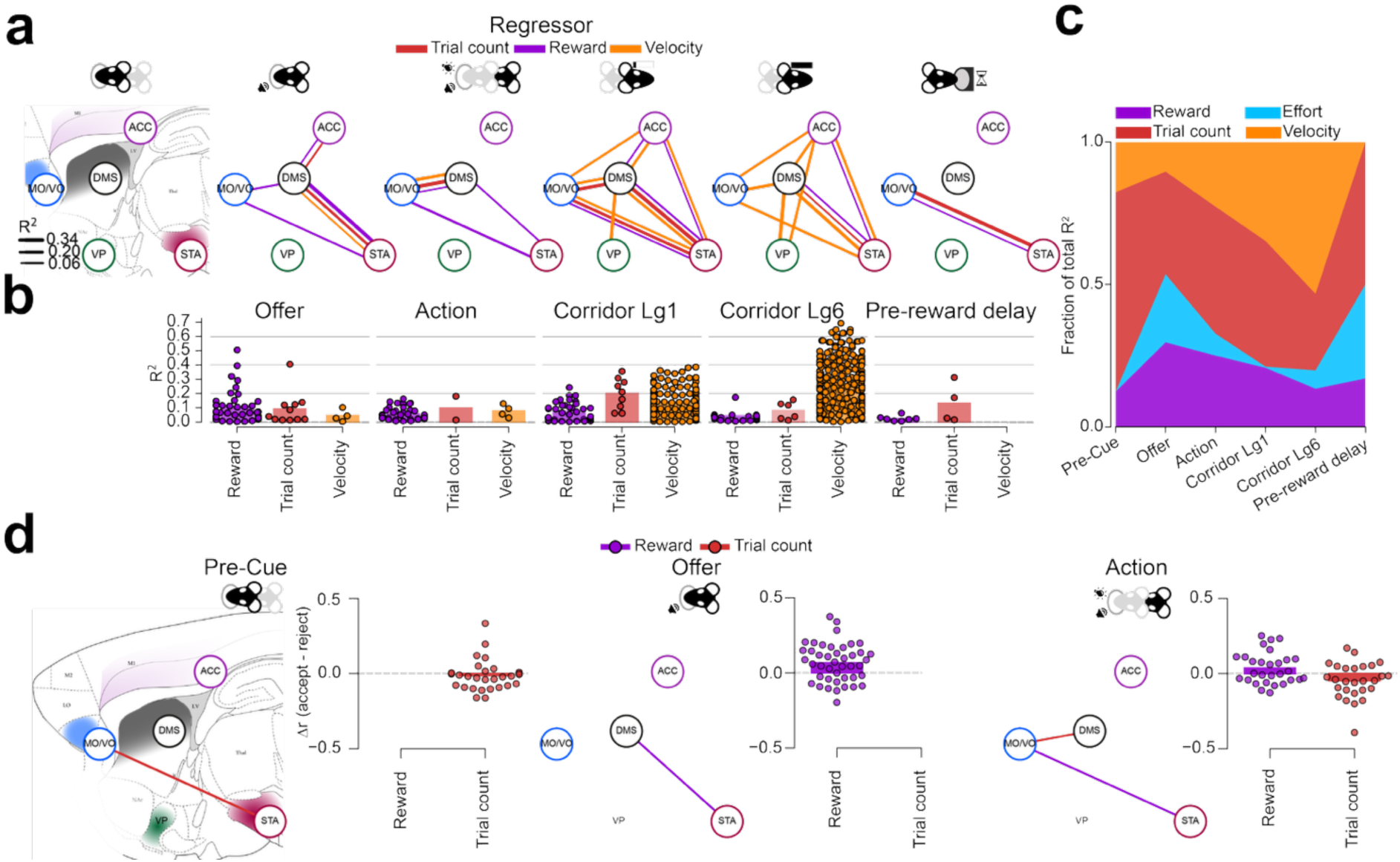
Shared subspaces in frontal-basal ganglia area pairs represent decision variables with temporal and spatial specificity gated by motivation (n = 68 sessions from 4 subjects). **a,** Network graphs showing significant decoding edges between area pairs for each trial epoch. An edge is drawn between two areas when a linear decoder trained on their shared CCA subspace predicted a given task variable (reward order, effort level, trial count, or velocity) significantly better than the permutated null data (one-tailed Wilcoxon signed-rank test on real vs. permuted cross-validated R², paired across subject-days; α = .01, uncorrected, as a BH correction was already applied on the CCA dimensions contributing to these decoders). Edge colour denotes the decoded task variable; edge width is scaled to the median out-of-sample R² across subject-days. **b,** Cross-validated decoding R² for each task variable, per epoch, restricted to area pairs significant in (a) and pooled across recording sessions. Bars show the across-session mean; swarm-plotted dots show individual significant session fits contributing to panel (a) and are colour-matched to panel a. Epochs with no significant edges are shown as empty. **c,** stacked composition of decoding R² across epochs. For each epoch, the fraction of total (positive) R² attributable to each task variable, averaged across significant area pairs and sessions. R² throughout is the out-of-sample coefficient of determination (1 − SSE/SST) from 10-fold cross-validated linear regression of each task variable on the significant population CC-subspace activity (k ≤ 10 components), with statistical significance assessed against 1,000 stage-shuffled permutations. **d,** Network diagrams showing, for each of the three epochs preceding reward pursuit (Pre-Cue, Offer, Action) whether that region pair’s held-out decoding of a task variable from the shared canonical-correlate subspace differed reliably between accepted and rejected trials. An edge is drawn only where the accept − reject difference in held-out decoding correlation (Δr) was both individually significant for that area pair (two-sided Wilcoxon signed-rank test of the per-session Δcorr against zero, p < .01) and survived a second, group-level test pooling every significant area pair for that task variable within the epoch (Wilcoxon vs. zero, Benjamini–Hochberg FDR-corrected across task variables, p < .01). Edges are coloured by task variable (reward, effort, trial count, running velocity; see legend); an area pair significant for more than one task variable is drawn as parallel offset lines. Line width is fixed and does not encode effect size. Insets depict the session-level Δ*r* (Pearson *r* between the true and decoded outcome variable, with the difference depicted between held-out accept and reject trial datasets), pooled across every area pair contributing a significant edge in panel **a.** Bars show the across-session mean; points are individual recording sessions.

Having established that shared cross-area activity carries task-relevant information during accepted trials, we next asked whether the these depended on the animal’s subsequent choice. We hypothesised that incentives like reward will be more strongly represented in the shared latent activity of the network on accepted trials, while travel cost (effort) and fatigue (trial count) may dominate this activity on rejected trials. We thus compared decoder performance between the two trial outcomes across equivalent trial segments (pre-cue, offer, action). For each region pair, we trained a decoder within the shared canonical correlation subspace — identified from data spanning both accepted and rejected trials — to predict each task variable (reward, effort, trial count, velocity). We then compared its held-out decoding accuracy (Pearson r between ground truth and predicted outcome), calculated on accepted or rejected trials in isolation. This revealed several significant differences between task variable decoding strength (fig. 3d). Most notably, reward representation was stronger in DMS–STA and MO/VO–STA canonical correlation subspaces during the offer and action epochs, respectively, on accepted trials. By contrast, trial count was more reliably decoded from MO/VO-STA shared activity during the pre-cue window of rejected trials.

To test whether these coding differences persisted after the decision-making window, we focused on the two corridor pursuit epochs in accepted trials and temporally equivalent trial segments on rejected trials. As the corridor was traversed on only 3.47% of rejected trials, an equal portion of time around corridor light-gate 1 and 6 crossings was extracted with the same latency offset from Go cue as in accepted trials. This revealed that stronger representation of reward in the frontal-basal ganglia and intrinsic basal ganglia network persisted during pursuit epochs (fig. S3c). Higher reward decoding performance on accepted trials was not driven by higher reward offer variance on those trials (fig. S3d). Together, these results suggest that the frontal-basal ganglia network transitions between distinct information processing states where the informational content reflects the animal’s underlying motivation to pursue an offer.

### Cross-area coupling is linked to individual differences in reward and effort sensitivity

Having established that shared cross-area activity represents decision variables differently depending on subsequent choice, we next asked whether the strength of inter-area coupling was associated with the degree to which animals’ choices were sensitive to reward or effort. Given the enhanced representation of reward in MO/VO- and DMS-STA activity on accepted trials, and of trial count in MO/VO–DMS activity on rejected trials, we asked whether the dominant canonical correlation in these circuits covaried with behavioural coefficient loadings within animal. To this end, we extracted the slopes of reward, effort and trial count on choices from generalised linear models fitted on each day’s behavioural data (fig. 4a, e; fig. S4a). Next, we regressed these on the z-scored canonical correlation of the first components (ρ_1_) from each area pair across all available sessions and animals by linear mixed effect modelling with a per-subject intercept. Surprisingly, this revealed that higher DMS-STA coupling along the first canonical correlation dimension (ρ_1_) was *negatively* associated with reward sensitivity within a given animal (fig. 4b-d; β = −0.11 per SD of ρ_1_, 95% CI −0.20 to −0.03, p = .009). By contrast, higher MO/VO-DMS ρ_1_ was associated with less within-animal dependence of choice on effort (fig. 4f-h; β = +0.13 per SD, 95% CI +0.05 to +0.21, p = .001). Sensitivity to trial count was not meaningfully related to ρ_1_ in any area pair (fig. S4). Taken together, cross-area coupling is reliably linked to behavioural sensitivity to reward and effort.

**Fig 4.**
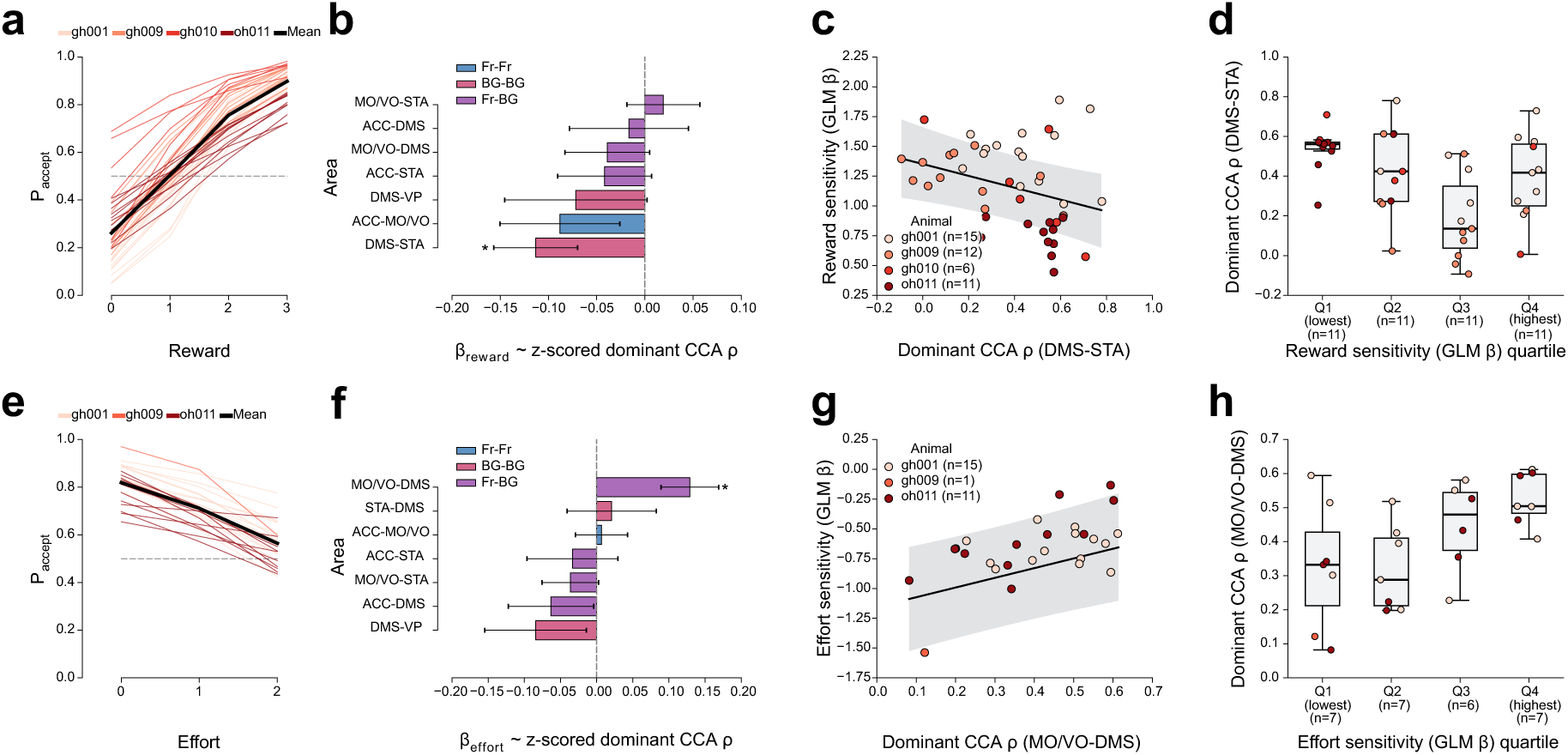
Cross-area coupling is associated with reward and effort sensitivity. **a, e,** Fitted probability of accepting an offer against reward (**a**) or effort (**e**) for each session (thin lines, coloured by animal) and the cohort mean (thick black line). Curves come from per-session binomial GLMs of accept/reject choice on reward, effort and cumulative reward, evaluated at the levels observed in that session with the other two regressors at the session median. Sessions shown are those in **c,d** (**a**; n = 44) and **g,h** (**e**; n = 27). **b, f,** Slopes of reward (**b**) or effort **(f)** sensitivity on the session’s dominant CCA ρ (ρ_1_), for each area pair. ρ is the held-out (10-fold cross-validated) first canonical correlation over the decision window (STAR methods), z-scored within area pair, so β is the change in sensitivity per 1 SD of the dominant ρ_1_ signal. Slopes are from linear mixed models with a random intercept for animal. Bars show β ± S.E.M and are coloured by circuit class (Fr–Fr, BG–BG, Fr–BG). *, uncorrected p < .01. **c, g,** Dominant ρ against reward sensitivity for STA–DMS and against effort sensitivity for MO/VO–DMS. Dots are sessions coloured by animal; the line and shading are the population-level fit and 95% CI of the mixed model in **b,f**. **d, h,** ρ_1_ across quartiles of reward sensitivity for STA–DMS (**d**; Kruskal-Wallis p = .005) and of effort sensitivity for MO/VO–DMS (**h;** Kruskal-Wallis p = .04), with sessions pooled across animals. Effort coefficients are negative, so Q1 holds the most effort-averse sessions. Boxes show the median and interquartile range, whiskers extend to 1.5 × the interquartile range, and dots are sessions coloured by animal.

## Discussion

Our results suggest that cost–benefit decision-making and subsequent reward pursuit are accompanied by a dynamic reconfiguration of communication across the frontal–basal ganglia network. Shared activity channels were found across frontal–frontal, frontal–basal ganglia and intrinsic basal ganglia connections, and their dimensionality varied markedly across task epochs and circuit classes. In particular, the number of significant shared dimensions increased within frontal–basal ganglia and intrinsic basal ganglia circuits during decision-making and reward pursuit, indicating that cross-area coupling is not a static property of the network but is recruited in a behaviourally dependent manner. Previous work has demonstrated that high-dimensional population representations, arising from heterogeneous mixed selectivity, can increase the computational flexibility of neural circuits by enabling a greater repertoire of input–output transformations (Fusi et al., 2016; Rigotti et al., 2013). In this context, the expansion of shared frontal–basal ganglia dimensionality may reflect the recruitment of a more diverse set of coordinated population patterns to support the multiple behavioural demands of cost-benefit decision-making and reward pursuit. Overall, these findings support the view that distributed decision-related activity is organised through transient, functionally distinct communication channels between interconnected frontal-basal ganglia structures.

The dimensionality of these shared representations also differed markedly between circuit classes. Frontal–basal ganglia interactions exhibited the most prominent expansion in dimensionality, particularly during reward pursuit, whereas intrinsic basal ganglia interactions exhibited a more modest number of significant dimensions. By contrast, frontal-frontal signal dimensionality was mostly one dimensional. These differences were not driven by disparities between recorded neuron counts in these circuit sub-classes. One possibility is that this organisation reflects the underlying anatomical architecture, including the substantial convergence of frontal cortical projections onto striatal neurons, with roughly 10 pyramidal neurons per striatal project neuron (Zheng & Wilson, 2002). Such convergence provides an anatomical substrate for the integration of multiple concurrent signals, consistent with the role of basal ganglia in transforming task state and motivational information into behavioural output (Haber, 2016; Klaus et al., 2019; Sharpe et al., 2019). Previous observations of close firing-rate covariation between frontal cortical areas and their topographically connected striatal targets during sensorimotor behaviour support the idea that these pathways exhibit coordinated activity (Peters et al., 2021). Our results extend this work by demonstrating that the dimensionality of this shared activity depends on the cognitive demands of the task. Together, these observations suggest that the organisation of shared population activity reflects an interaction between the anatomical architecture of the network and the demands of ongoing behaviour.

The information carried by these shared dimensions was similarly structured in time and across circuit classes. Before offer cue onset, we found no evidence that shared activity encoded the task variables, providing an important baseline against which subsequent coding could be interpreted. Following offer presentation, reward information became prominently represented within frontal–basal ganglia interactions and, to a lesser extent, intrinsic basal ganglia interactions. Reward-related coding persisted into periods of reward pursuit, but was increasingly accompanied, and ultimately superseded, by representations of instantaneous velocity. Although our analyses cannot exclude the possibility that this transition partly reflects the greater variance in velocity during corridor pursuit, the overall pattern is consistent with a shift in shared population activity from representing prospective outcomes towards coordinating the behavioural processes required to obtain them. Trial count, which may capture slowly evolving factors such as time on task, satiety, fatigue or motivational state, was also represented within several shared frontal–basal ganglia dimensions, particularly during offer presentation. Thus, cross-area coupling appears to simultaneously carry both externally defined task variables and internally evolving behavioural-state information. Importantly, these signals were not distributed indiscriminately across the network: reward representations were preferentially expressed in frontal–basal ganglia and some intrinsic basal ganglia interactions, whereas velocity-related information was considerably more widespread. This circuit specificity supports the idea that distinct frontal–basal ganglia motifs make different contributions to decision formation and behavioural implementation.

Although effort-related firing has been reported in individual neurons and co-active ensembles populations within several components of the frontal–basal ganglia network (Härmson et al, 2025), effort could not be significantly decoded from the low-dimensional shared activity identified here. This may be a reflection of the modest underlying effect size of single unit representations of effort (Härmson et al., 2025). However, the dissociation may also suggest that information represented within individual regions need not necessarily be transmitted in the dominant dimensions of activity shared between them. Effort may instead be encoded predominantly within local or non-shared population dimensions, communicated through weaker or higher-dimensional interactions not captured by the present CCA approach with the observed cell yields, or represented in a temporally specific form that is not well suited for linear regression detection across trials in session. Our findings do not preclude the presence of effort representation in the shared low-dimensional activity of areas such as the cingulate motor cortex or dorsolateral prefrontal cortex (Hosokawa et al., 2013), nor do they preclude more prominent representation of effort when this is varied trial-by-trial (Hillman & Bilkey, 2010; Porter et al., 2019). Nonetheless, our findings show that parallel representations of a variable across anatomically connected areas need not necessarily reflect direct communication of that variable between them. Shared latent activity appears to constitute a selective subset of the information available within each region.

The dependence of shared information on subsequent choice provides further evidence that cross-area communication is dynamically organised according to behavioural state. Reward was more strongly represented within frontal–basal ganglia and intrinsic basal ganglia interactions *on accepted trials*, particularly during offer evaluation and pursuit. By contrast, trial count was more strongly represented on *rejected trials* during trial initiation as well as during time-matched periods of corridor running, suggesting that the network placed greater weight on slowly evolving changes in internal state such as increasing fatigue or satiety. This selective representation may be driven by a state-dependent reconfiguration of the latent space which prioritises certain decision variables over others depending on current physiological needs. Future work will be needed to elaborate whether there is neural evidence of attractor states that can explain this selective re-weighting of representations. Nevertheless, the overall pattern suggests that frontal–basal ganglia communication does not operate in a fixed representational configuration; rather, the reward-, state- and movement-related information are selectively re-weighted depending on the underlying motivation to pursue an offer.

Intriguingly, cross-area coupling was not only associated with decision-related trial-by-trial information, but also how subjects weighed costs and benefits to make decisions over sessions. Within a given animal, higher DMS-STA coupling was associated with reduced reward sensitivity. By contrast, stronger MO/VO-DMS coupling was associated with less negative effort coefficients, indicative of reduced effort aversion. The absence of a detectable effort signal in the decoding analysis need not conflict with these findings. Linear decoders assume a stable sign-consistent readout within each session, so effort effects that vary in sign over time could go undetected, whereas relating coupling to session-level behavioural coefficients does not depend on such a readout. These findings raise the possibility that motivational deficits may arise from dysfunction in specific communication pathways that differentially influence the valuation of prospective rewards and the willingness to expend effort. Supporting this possibility, a recent clinical study linked reduced structural connectivity between the dorsal anterior cingulate cortex and striatum to a greater tendency to reject high-effort offers in patients with Parkinson’s disease and apathy (Morris et al., 2026). Similarly, anhedonia severity in major depressive disorder has been associated with reduced task-dependent functional connectivity between the posterior ventromedial prefrontal cortex and interconnected reward-processing regions, including the nucleus accumbens and orbitofrontal cortex during the processing of pleasant stimuli (Young et al., 2016). This circuit- and context-specific perspective may have translational relevance for the development of targeted interventions to ameliorate motivational symptoms, by suggesting that effective therapies may need to modulate distinct neural pathways according to the particular component of motivated behaviour they seek to restore. However, establishing whether these pathways causally regulate reward and effort sensitivity, and whether their modulation can ameliorate motivational deficits, will require further investigation.

Taken together, our findings support a circuit-specific account of frontal–basal ganglia communication during cost–benefit behaviour. Decision formation recruited frontal–basal ganglia communication channels carrying prospective reward information alongside signals associated with slower changes in behavioural state, whereas reward pursuit increasingly engaged widespread representations of movement. Intrinsic basal ganglia coupling contributed to both reward-related and movement coding, consistent with a role in linking motivational information to the execution of goal-directed action. The information carried by these circuits was also choice-dependent: reward was more strongly represented on accepted trials, whereas trial count was more strongly represented on rejected trials, consistent with a greater influence of evolving internal state on decisions to forgo an offer. Moreover, coupling strength between frontal-basal ganglia and intrinsic basal ganglia areas was associated with individual differences in behavioural sensitivity. Our findings extend previous accounts of distributed neural coding and suggest that cost–benefit behaviour is paralleled by dynamic recruitment of subcircuits carrying spatiotemporally specific information about prospective reward, behavioural state and ongoing action. These circuit-specific associations provide a basis for investigating whether targeted modulation of distinct communication pathways could help address different components of motivational dysfunction.

### Limitations of the study

This study has several limitations relating to the estimation of shared population dimensionality, their consistency between animals, and its causal relation to cost-benefit decision making and reward pursuit. First, the dimensionality of shared activity identified here may depend partly on the number of simultaneously recorded neurons and available trials. Although differences in dimensionality were not readily explained by variation in circuit class unit yield, it remains possible that additional shared dimensions would emerge with greater sampling of neuronal populations or trial numbers. Thus, the dimensions identified here should be considered an estimate of statistically detectable shared population activity, rather than necessarily representing the full dimensionality of inter-area interactions. Second, some area pairs, notably MO/VO–ACC and MO/VO–VP, were recorded in only a few sessions from a small number of animals, precluding reliable assessment of between-animal consistency. Estimates for these pairs should thus be interpreted with caution. Third, CCA identifies shared patterns of correlated population activity but does not establish the direction of information transfer or distinguish direct communication from shared inputs or other sources of correlated activity. Accordingly, the recruitment of shared activity channels and their association with behavioural sensitivity do not, in themselves, demonstrate a causal role for these dynamics in cost–benefit decision-making or reward pursuit. Future studies combining more extensive and balanced neuronal sampling across animals with causal manipulation of specific pathways will be important for establishing the extent of shared population dimensionality, its generalisability across individuals, and its causal contribution to motivated behaviour.

### Resource Availability

Lead contact: Andrew Sharott

- Data Availability: Data is available at 10.60964/rnd-1m60-zh48.
- Code Availability: Analysis code used in this study is available at https://github.com/oliverharmson/costbenefit-cca.git.
- Materials Availability: The study did not use any unique reagents or materials.

Further information and requests for resources and reagents should be directed to and will be fulfilled by the Lead Contact, Andrew Sharott.

## Acknowledgements

We would like to thank Ben Micklem for methodological and technical assistance during the project. This work was supported by The Clarendon Fund and Mary Somerville Clarendon Graduate Scholarship in conjunction with the Department of Experimental Psychology (award SFF1819_CB2_MSD_ 1196514 to OH) and the Medical Research Council UK (awards MC_UU_00003/6 and UKRI/MR/B000936/1 to AS; award MR/X022080/1 to MW, AS and SM).

## Author contributions

Conceptualisation: OH, SGM, MEW and AS; Technical development: OH, RT, BP; Investigation: OH, BP and IG; Analysis: OH; Resources and funding acquisition: AS, MEW and SGM; Writing - original draft: OH; Writing - Reviewing & Editing: OH, IG, BP, SGM, MEW, AS; Supervision: AS, SGM, MEW.

## STAR Methods

### EXPERIMENTAL MODEL AND STUDY PARTICIPANT DETAILS

All procedures were carried out in accordance with the UK Animals (Scientific Procedures) Act (1986) and its associated guidelines. We used 12 group-housed Lister Hooded naive male rats (Charles River, U.K.) weighing 300-390g at the beginning of training. Rats were maintained on a twelve-hour light/dark cycle (lights on at 7 AM). During testing periods, rats were food restricted to a target weight range of 85-90% of their free-feeding weight. Water was available ad libitum in their home cages. Following implantation with multielectrode drives, animals were singly housed.

### METHOD DETAILS

All listed analyses are implemented on data from a prior publication focused on single neuron and co-active ensemble analyses (Härmson et al., 2025). Here, we describe novel methods related to analyses of neurons’ shared activity in pairs of areas. While we briefly describe previously published methodological details here for accessibility, we refer readers to the original work in-text.

#### Experimental subject details

All procedures were carried out in accordance with the UK Animals (Scientific Procedures) Act (1986) and its associated guidelines. We used 12 group-housed Lister Hooded naive rats (Charles River, U.K.) weighing 300-390g at the beginning of training. Rats were maintained on a twelve-hour light/dark cycle (lights on at 7 AM). During testing periods, rats were food restricted to a target weight range of 85-90% of their free-feeding weight. Water was available *ad libitum* in their home cages. Following implantation with multielectrode drives, animals were singly housed.

#### Behavioural apparatus

The operant box consisted of two 17.6cm diameter custom-designed 3D-printed plastic semi-circular arenas, connected by a 130 cm-long Perspex corridor with walls on either side (Härmson et al., 2025). The corridor had 6 equidistantly placed infra-red (IR) light-gates to register the rats’ position and 3 equidistant slots for barriers to increase the travel cost between the ends of the corridor. Each arena housed a custom designed 3D-printed plastic nose-poke and reward port, positioned symmetrically on either side. Nose-poke and reward ports contained a infrared light-gate and a white LED house light. Reward ports were connected with 20-mg food pellet dispensers (Med-associates) fitted outside the walls of the arenas using medical-grade silicone tubing. The electronics were controlled by a Teensy 3.6 microcontroller that ran on custom-written firmware. The apparatus was lit using far-red LEDs to minimise visibility during training and testing.

#### Behavioural paradigm and training

Rats were handled for a week prior to training. Training consisted of 5 steps, described in detail in (Härmson et al., 2025). Rats initiated trials by poking and holding in the nose-poke port of the starting arena (“pre-cue”). The starting arena was randomly allocated on the 1st trial. After a pre-cue period (0.3-0.4s), an auditory cue (“offer”) was again presented from the distal end of the apparatus to signal the available reward at end of the track from where they initiated. The available reward was pseudo-randomly varied trial-by-trial (4 sizes: nil, low, mid, high, corresponding to zero, one, three or six 20mg sucrose pellets), signalled by a distinct auditory cue. The cue-reward associations were counter-balanced across subjects. Rats were required to maintain nose-poking for another 0.6-0.8s until the onset of a visual Go cue, signalled by a white light LED. Premature exiting of the nose-poke either during the pre-cue or offer period would elicit a 5-s error time-out, indicated by a flashing house-light in the nose-poke, followed by a 5s ITI. If rats exited prematurely during the auditory cue, the same cue would be repeated on the next nose-poke attempt. A grace period of up to 100ms during nose-poking - where nose-poke exits did not count towards an error - allowed for minor adjustments in body position. To accept the offer, rats needed to enter the corridor (cross a light-gate 24cm away from the arena) within a time window; otherwise, that offer was rejected. The cost of pursuing reward was pseudo-randomly varied across blocks of 17-23 trials (3 levels: low, moderate, high), implemented using varying numbers of barriers (0-2 or 1-3 in implanted or non-implanted animals, respectively) in the corridor. The barriers were 0.5 cm thick and were 12 cm and 18 cm tall at their lowest and highest points respectively, with a 45-degree slant mid-way between to generate a more effortful zig-zag of running between the ends of the corridor.

#### Multielectrode drive

For single unit recordings we used a bespoke driveable multielectrode device weighing ∼20g. The implant was comprised of a lid which housed a 128-pin connector designed for Intan 64-channel head-stages (Intan Technology), a plastic corpus that housed the electrode movement mechanisms, glass tubing and a base. All components were assembled in house and the assembly steps are described in detail in (Härmson et al., 2025).

Three of the subjects (gh001, gh009, gh010) reported here received an implant that targeted all 5 regions, whereas one subject (oh011) received a modified drive without the VP electrodes (table 1). Electrodes in the MO/VO, ACC, and VP were arranged as a row along one sagittal plane. DMS electrodes were as rows along two sagittal planes and STA electrodes were arranged in triangular fashion around a centre point. The ML and AP positions of each electrode in the drive were planned around a centre-point co-ordinate obtained as an average between readings in two different versions of a commonly used rat brain atlas (Paxinos & Watson, 1998, 2006) that visually offered the most exposure in each structure, while also avoiding the ventricle. Relative electrode positions in the AP and ML dimensions were planned around centroid co-ordinates.

#### Implantation of multielectrode drives for recordings in behaving rats

Electrode implantations were performed in a subset of expert animals (n=4; table 1). Surgeries were performed under isoflurane anaesthesia (5% for induction, 1.5-2.5% maintenance) and oxygen (2L/min). Subcutaneous local anaesthetic (Marcaine, 2mg/kg, 2.5 mg/mL) and a non-steroidal anti-inflammatory drug (Metacam, 1mg/kg, 5mg/mL) were administered at the beginning of each surgery. Post-operative opioid analgesia (Vetergersic, 0.3 mg/mL, 0.03 mg/kg) was administered subcutaneously for three consecutive post-operative days.

Craniotomies were created above each of the targeted areas and *dura mater* thoroughly removed from the visible brain surface. The drive was aligned on the animals’ heads by using the centre-point of the STA electrode bundle as a reference. Next, the drive was lowered to target depth on a stereotaxic frame (David Kopf Instruments) based on the DV reading of the STA electrode bundle from brain surface. The implant was then secured to skull-attached screws using adhesive dental cement and dental acrylic. Recordings were referenced to two supra-cerebellar screws which were connected to the implant during surgery. Neuronal recordings began at least 2 weeks after surgery. Electrode placement was confirmed using histology.

#### Histological data analysis

To verify electrode placements, after the experiments were completed, animals were deeply anaesthetised with isoflurane (5%) and intraperitoneal pentobarbital (3 mL, Pentoject, 200 mg/mL and perfused trans-cardially with phosphate buffered saline (PBS) followed by fixative solution (paraformaldehyde dissolved in PBS, 4%, wt/vol). For post-mortem fixation, the sectioned heads were further submerged in fixative for another 24h, following which brains were extracted and washed 3 times in PBS and stored in a PBS solution containing 0.02% sodium azide until sectioning. Brains were sectioned at 0.1mm thickness using a vibratome (Leica), immunohistologically stained for FoxP2 and TH, and mounted in Vectashield. Unfiltered “bright-field” epifluorescence microscope images were acquired using 5x magnification and Zen Blue Pro (Zeiss).

#### Electrode localisation and histological verification

To estimate all the possible recorded areas (fig. S1) and exclude cells yields from neighbouring regions, we (1) estimated each identifiable electrode tract’s DV end-depth from our epifluorescence imaging data, (2) averaged this across all identifiable electrode tracts to get an average end-depth, (3) estimated each day’s targeted structure based on a log of putative lowering distances averaged across electrodes within a region. We did not restrict the cell yield for the STA based on histological data. For other structures we restricted our cell yields to only the MO or VO for MO/VO, Cg1 and Cg2 for the ACC, CPu for the DMS; and VP only for the VP. Estimation of the position of the electrode tip on a given recording day was based on the estimated extent of electrode advancement (i.e. the number of turns of the screw driving the electrode within the implant) and the tract location in the histological data. Wherever neurons were detected on days where the anatomical localisation of the electrode tip suggested we were either out of the brain near a target structure or in a nearby white matter tract, but that electrode trajectory was consistent with being in the target areas, we presumed that the error was in the estimate of the electrode tip and these cells were included in the analysed cell yield for that area. Neurons were not included if the entire electrode trajectory was not on target.

#### Electrophysiology data acquisition and processing

Wideband and accelerometer data were recorded, amplified and digitised on 2 tethered Intan 64-channel head-stages at 20,000 Hz using an Intan acquisition board (Intan Technologies). No hardware filtering was applied during recording. Files were saved in binary format. Each electrode was lowered by at least 125 μm up to 2h ahead of each recording session to ensure identification of unique units every day.

#### Spike sorting

Clustering and curation of units were carried out in KiloSort2(Marius Pachitariu, 2016) and Phy (available at https://phy.readthedocs.io/en/latest/), respectively. Both are open-source libraries optimised for high-density multielectrode recordings. Individual units were isolated from the broadband signal based on waveforms using the KiloSort2 automatic clustering algorithm. A number of quality control steps were used in order to verify the single units identified by KiloSort2. Each neuron was manually inspected in Phy, and multiunits were excluded. Single units were first screened by the shape of the waveform. Only neurons with a stable waveform with spikes present across the entire recording block were included in further analysis. Next, we inspected the ISIs of each single unit to ensure that a clear refractory period was present. Cluster cutting was used to remove any contamination by electrical artefacts. Finally, the cross correlation between single units and their waveforms were used to merge those single units with a matching waveform and shared refractory period. Finally, single units with a firing rate less than <0.1Hz were excluded. No multi-units were used in the reported set of analyses.

#### DeepLabCut Video Analyses

We carried out the above analysis on the positional data extracted from DeepLabCut. We trained the resnet_50 model on labelled frames from 664 videos (3 frames labelled per video) which spanned all four recorded animals across all configurations of barriers. Specifically, we labelled the position of the back of the head of animals, between their ears. The model was trained over 900,000 iterations until the loss had plateaued. Model performance was then checked by visual inspection of labelled videos and using the alignment with light gate data The results and verification of this have been described previously (Härmson et al., 2025).

### Neuronal population analyses

#### Firing rate matrix construction

Spike times were binned in 25 ms windows and firing rates were smoothed with a causal exponential smoothing filter (constant τ = 50ms and a kernel radius of 200ms). These data were reshaped into trial × unit arrays per epoch, session, and animal. For each ordered pair of the five regions, only session-days on which at least one unit was recorded simultaneously in both regions were retained, yielding a per-session-day pool of qualifying unit pairs for cross-area analysis.

For visualising task event-locked firing rates for representative neurons (Fig. S1a), spiking counts were quantified in 25ms *non-overlapping* bins sampled -400ms to +400ms around the initiation of the nose-poke (termed “Pre-Cue”), -400ms to +400ms around the onset of the offer cue (termed “Offer”), -500ms to +500ms around the onset of the Go cue (termed “Go Cue”) and -500ms pre-to +1000ms following nose-poke exit (termed “Action”). Data were subsequently averaged across trials of the same magnitude of offered reward, the required effort and prospective decision, each time averaging across the other two variables. Peri-stimulus time histogram (PSTH) data were smoothed with a causal exponential smoothing filter with the same parameters as above.

#### Cross-validated canonical correlation analysis (*fig. 1*-2 and S1-2)

Cortical–basal ganglia communication was quantified with cross-validated canonical correlation analysis (CCA) applied to simultaneously recorded single units (rate ≥ 0.1 Hz) from MO/VO, ACC, DMS, VP and STA in [4] rats over [14] sessions. Spikes were binned at 25 ms, smoothed with a causal exponential kernel (τ = 50 ms), and averaged within each of six task epochs (Pre-cue, Offer, Action, Corridor Lg1, Corridor Lg6 and Pre-reward delay). This gave one firing-rate value per unit, trial and epoch. For every session, area pair and epoch, up to ten canonical components were fitted (scikit-learn CCA, standardised columns; units active in fewer than 20% of trials were excluded). Weights were estimated on nine of ten contiguous trial blocks, and the Pearson correlation (ρ) between the two areas’ canonical variates was computed on the held-out block. Fold-wise ρ values were averaged with the Fisher z-transformation. A null distribution was generated by randomly shuffling one area’s held-out activity across stages within a recording block (1,000 shifts), which preserves temporal autocorrelation and underlying staged structure of each recording block, but breaks cross-area trial correspondence. Edges in the network diagrams (Fig. 2a) were retained where the observed ρ exceeded the mean permuted ρ across sessions (one-tailed Wilcoxon signed-rank test, Benjamini–Hochberg FDR = .01, pooled across epochs, area pairs and component ranks). Edge weights are session-averaged Δρ (|ρ*_observed_| − |ρ*_permuted_|). Variability in Δρ was estimated separately for each epoch and circuit class (Fr–Fr, Fr–BG, BG–BG) by pooling session-level values from significant edges and bootstrapping the mean (5,000 resamples, percentile 95% CI).

Communication dimensionality (Fig. 2b) was defined as the number of significant canonical components. Components were counted per session, epoch and circuit class, using per-session permutation p-values corrected within epoch (Benjamini–Hochberg, FDR = .01), and summed over the area pairs in each class. Counts were analysed with a two-way repeated-measures ANOVA (circuit class × epoch, session as the repeated measure; sessions with any missing cell were excluded; [Greenhouse–Geisser-corrected p-values]). Effect sizes are generalised eta-squared. Post-hoc paired t-tests were run within each family of comparisons (circuit class, epoch and circuit × epoch cells), corrected with Benjamini–Hochberg (FDR = .01), and effect sizes are reported as Hedges’ g.

Because the number of components that can reach significance depends on how many neurons were recorded, we tested whether unit yield differed between circuit classes. Daily yield was defined as the number of units retained after the firing-rate criterion in each area on each recording day, with each unit counted once per day even if it was recorded across several blocks. The combined yield of a pair was the sum of the two areas’ yields for days on which both areas were recorded. This formed the population input for CCA models. Combined yields were log-transformed, so that effects are ratios, and modelled with a linear mixed model with circuit class as a fixed effect, a random intercept for animal, and a variance component for recording day nested within animal, to account for days contributing to several pairs and for clustering within animals. The effect of circuit class was assessed with a likelihood-ratio test against an intercept-only model (both fitted by maximum likelihood), and post-hoc contrasts used Wald tests on the restricted-maximum-likelihood fit with Bonferroni correction.

Because the number of components that can reach significance depends on how many neurons were recorded, we tested whether unit yield differed between circuit classes. Daily yield was defined as the number of units retained after the firing-rate criterion in each area on each recording day, with each unit counted once per day even if it was recorded across several blocks. The combined yield of a pair was the sum of the two areas’ yields, for days on which both areas were recorded; this is the population entering that pair’s CCA. Combined yields were log-transformed, so that effects are ratios, and modelled with a linear mixed model with circuit class as a fixed effect, a random intercept for animal, and a variance component for recording day nested within animal, to account for days contributing to several pairs and for clustering within animals. The effect of circuit class was assessed with a likelihood-ratio test against an intercept-only model (both fitted by maximum likelihood), and post-hoc contrasts used Wald tests on the restricted-maximum-likelihood fit with Bonferroni correction.

Between-animal reliability of the region-pair correlation estimates (fig. S1) was quantified as an intraclass correlation coefficient (ICC), computed from a random-intercept linear mixed model (y ∼ 1 + (1|subject), fit with statsmodels mixedlm) as the ratio of between-subject to within-subject (residual) variance; lower ICC indicates greater cross-animal agreement. 95% confidence intervals were obtained via a subject-stratified bootstrap (5000 resamples, resampling sessions with replacement within each animal).

#### Decoding task variables from cross-validated CCA subspaces (*fig. 3* and S3)

To ask which task variables are represented by the shared cortical–basal ganglia subspaces identified in Fig. 2, we regressed trial-wise task variables onto canonical component (CC) activity. Trials were divided into 10 contiguous blocks, and a CCA was fitted on the trials of nine blocks and used to project the trials of the held-out block. This yielded an out-of-fold CC score for every trial, so that no trial contributed to the weights that produced its own score. We modelled every session × area pair × epoch that had at least one significant canonical dimension (Fig. 2a). For each of the two areas in a pair, one task variable at a time (reward offer, effort level, trial number, and running speed) was decoded from that area’s CC scores. The target was the trial’s value of the variable, and the predictors were all (up to 10) CC scores. Running speed was computed from DeepLabCut-tracked position, averaged over each epoch window. Models were ordinary least-squares regressions with z-scored predictors and target, fitted within 10-fold contiguous-block cross-validation. Out-of-fold predictions were pooled and scored as R². Scores were then averaged across the two regions of each analysed area pair within each session. Significance was assessed with a permutation test (1,000 permutations). The p-value was defined as (b + 1)/(n + 1). Only models with R² above zero and above the permuted mean were carried forward. For each task variable, epoch and area pair we then tested whether session-level R² exceeded the permuted R² (one-tailed Wilcoxon signed-rank test, p < .01). Significant edges were plotted in Fig. 3a. Line colour indicates the task variable and line width the median R² across sessions. Fig. 3b shows per-session R² (bars: mean; dots: individual sessions) pooled across the significant area pairs for each task variable and epoch. Fig. 3c shows the fractional composition of R² across epochs, computed from the mean R² of each task variable across models with a permutation p < .01. Models with a negative R² were discarded and each epoch normalised to sum to 1.

#### Cross-epoch geometry of task variable decoding axes (Fig. S3a)

To test whether the variable that takes over from reward decoding uses the same neural direction as earlier reward coding, we compared decoding axes across epochs in the space of the recorded units. CCA models were fitted as described above. Reward, trial count, velocity and effort were projected onto the significant shared subspaces of each area pair as above. Because the CCA is fit separately in every epoch, each CCA component k is a different direction in each epoch, so fold-averaged coefficients cannot be compared across epochs. We therefore mapped each decoder back onto the units, u = W·b, where W holds the canonical weights of the standardised units. The cosine between the competing variable’s axis in a corridor epoch and the reward axis of an earlier epoch was then computed over the units present in both epochs’ analyses (≥5), separately for each area of the pair, and only where both variables were decodable. Session-level cosines were summarised by a linear mixed model (intercept only, REML; random intercept per animal and a session variance component nested within animal). We report the mean with a 98% interval and test it against zero (Wald). Equivalence to zero was tested against a ±0.2 margin (cos 0.2 = 78.5°; 4% shared squared length) using two one-sided tests (α = .01); both tests were Benjamini–Hochberg-corrected across the 24 comparisons shown.

#### Decoding task variables from shared cross-area activity on accepted and rejected trials (*fig. 3* and S3)

Shared activity between each pair of simultaneously recorded regions was defined by canonical correlation analysis (CCA) of binned, smoothed firing rates (25 ms bins), computed separately for each of five trial epochs (Pre-Cue, Offer, Action, and early and late corridor approach or equivalent time-matched periods in rejected trials) using data spanning both accepted and rejected trials. Canonical dimensions were retained for each area pair and epoch if they exceeded a permutation-based null (n = 1,000) in accepted or rejected trials. We then asked how well each pair’s retained canonical scores decode four task variables: reward magnitude, effort level, trial count and running velocity. Decoders were linear regressions trained with 5-fold cross-validation on trials pooled across both outcomes. A lower fold-validation number was used here to ensure sufficient training data was available given the relatively smaller amount of rejected trials. Held-out predictions were then scored separately for accepted and rejected trials as the Pearson correlation (*r*) between decoded and true values. Because the two decision outcomes differed in trial numbers, the larger group’s score in each fold was the Fisher-z average of r over B = 20 random subsamples, drawn without replacement, matched to the smaller group’s trial count.

For each session, area pair, epoch and task variable we computed Δ*r* = r_accept_ − r_reject_. An area pair was considered to show an outcome-dependent difference if its per-session Δ*r* values differed from zero (two-sided Wilcoxon signed-rank test), after Benjamini–Hochberg FDR correction across all epoch and area-pair tests for that variable (α = .01). Significant pairs are shown as network edges, coloured by task variable. Per-session Δr values from these pairs were then pooled for each variable and epoch for display. To test whether outcome differences in decoding could arise from differences in the spread of the decoded variables, we computed the variance of each variable’s true value within accepted and rejected trials for each session, applying the same subsampling procedure to the larger group (B = 20). We then compared the per-session log₂ variance ratio (accept/reject) against zero with a two-sided Wilcoxon signed-rank test (p < .01; fig. S4).

### Relating cortico-basal ganglia coupling to choice behaviour (Fig. 4, Fig. S4)

#### Behavioural model

Each session (a recording day) was fit separately with a binomial (logit) GLM, accept ∼ reward + effort + trial count, where trial count is the running trial index within the day (0 to ∼200). All 74 fits converged. Each fit yielded an intercept and a coefficient per regressor, which we refer to as the session’s reward, effort and trial-count sensitivity. Fitted choice curves (Fig. 4a,e; Fig. S4a) are the model’s P(accept) as a function of one regressor, evaluated at the levels observed in that session with the other two regressors held at the session’s median. Cohort means are the mean of the session curves at each level (for trial count, the mean of the interpolated curves up to the trial count reached by 75% of sessions).

#### Canonical correlation analysis

Neural coupling between simultaneously recorded areas (MO/VO, ACC, DMS, VP, STA) was quantified per session and area pair by canonical correlation analysis (CCA) of single-unit firing rates in the decision window as defined in (Härmson et al., 2025), ranging from the start of the pre-cue epoch to 900ms before observed corridor entry on accepted trials or 900ms before the per-session median of corridor entry latencies on rejected trials. For each trial, each unit’s firing rate (Hz) was the mean over this window, giving one trials × units matrix per area. Accepted and rejected trials were both included, error trials were excluded, units with mean firing rate below 0.1 Hz or active on fewer than 20% of a session’s trials were dropped, and at least two units per area were required. Canonical correlation analysis was performed as previously described.

#### Relating canonical correlation magnitude to behaviour

Sessions in which a pair had both a behavioural fit and a CCA result were used (13–49 sessions per pair, 2–4 animals). For each area pair, ρ_1_ was z-scored across its sessions, and each behavioural coefficient was regressed on it with a linear mixed model with a random intercept for animal (statsmodels MixedLM, coefficient scaled to unit variance for fitting and scaled back for reporting). Slopes are therefore changes in the coefficient per standard deviation of ρ. Only the first (dominant) canonical correlation dimension (ρ_1_) was analysed in this dataset. If only one animal contributed, or the fit was singular (animal variance estimated at zero; DMS–VP, trial count), an ordinary least squares fit was used. Quartile differences were tested with Kruskal–Wallis tests.

### ADDITIONAL RESOURCES

Data was analysed using the following open-source python packages/functions listed below (Key Resources’ Table).

#### Key resources table

Overview of all key code libraries used in this project.

| REAGENT or RESOURCE | SOURCE | IDENTIFIER |
|---|---|---|
| Deposited data |  |  |
| Spike timings: firing rates of all single units (25 ms bins), accepted and rejected trials | This paper | 10.60964/rnd-1m60-zh48 |
| Behavioural timestamps: event channels, behavioural states and trial tables | This paper | 10.60964/rnd-1m60-zh48 |
| Video tracking tables: DeepLabCut position of the animal (100 Hz) | This paper | 10.60964/rnd-1m60-zh48 |
| Cross-validated canonical correlation (CCA) and task-decoder results | This paper | 10.60964/rnd-1m60-zh48 |
| Software and algorithms |  |  |
| Analysis code (CCA dimensionality, task decoding, connectivity–behaviour and supplemental analyses) | This paper | https://github.com/oliverharmson/costbenefit-cca.git |
| Python 3.10: programming language | Python Software Foundation | https://www.python.org |
| stats from SciPy: wilcoxon() used for Wilcoxon signed-rank tests of canonical correlations against their permutation null and of accept–reject decoding differences against zero; kruskal() for the Kruskal–Wallis test and spearmanr() for Spearman correlations across sensitivity quartiles; friedmanchisquare() for the Friedman test of unit yields; ttest_1samp() for one-sample t-tests; chi2.sf() for likelihood-ratio tests; norm.sf(), norm.cdf() and norm.ppf() for Wald and equivalence tests and confidence limits; gaussian_kde() for kernel density estimates | (Harris et al., 2020) | https://docs.scipy.org/doc/scipy/reference/stats.html |
| ndimage from SciPy: gaussian_filter1d() used for optional Gaussian smoothing of binned firing rates | (Harris et al., 2020) | https://docs.scipy.org/doc/scipy/reference/ndimage.html |
| statsmodels.formula.api: mixedlm() used to fit linear mixed-effects models (intraclass correlations, unit yield, decoder-vector dot products, connectivity–behaviour regressions); glm() with the Binomial family for per-session logistic regression of accept/reject choice; ols() for linear regression | (Seabold & Perktold, 2010) | https://www.statsmodels.org/v0.14.5/ |
| statsmodels.stats.multitest: multipletests() used for multiple-comparison correction | (Seabold & Perktold, 2010) | https://www.statsmodels.org/v0.14.5/ |
| Pingouin: rm_anova() used for two-way repeated-measures ANOVA and pairwise_tests() for post-hoc comparisons | (Vallat, 2018) | https://pingouin-stats.org |
| NumPy: matrix/array operations; random.default_rng() with choice(), integers() and permutation() used for seeded bootstrap resampling, trial subsampling and null draws; linalg.norm() and arccos() used to compute angles between decoder weight vectors; percentile(), quantile() and histogram() for summary statistics | (Harris et al., 2020) | https://numpy.org |
| pandas: DataFrame() used for the storage and manipulation of data; groupby(), merge(), pivot(), qcut() and rolling() for aggregation, joining and binning; read_feather() and to_feather() for Feather tables | (MicKinney, 2010) | https://pandas.pydata.org |
| matplotlib.pyplot: ax.scatter(), ax.plot(), ax.bar(), ax.barh(), ax.errorbar(), ax.boxplot(), ax.fill_between(), ax.fill_betweenx(), ax.hist() and ax.imshow(), with gridspec, used for data visualisation | (Hunter, 2007) | https://matplotlib.org/3.8.4/api/ |
| seaborn: heatmap() used for the heatmaps of recordings per area pair and swarmplot() for session-level dot plots | (Waskom, 2021) | https://seaborn.pydata.org |
| transform from scikit-image: rescale() used to resize the epoch icons placed in figures | (van der Walt et al., 2014) | https://scikit-image.org |
| Ray: put(), remote(), get() and wait() used to parallelise the computation of firing-rate tables across CPU cores | (Moller & Bogacz, 2019) | https://www.ray.io |
| PyArrow: Apache Arrow/Feather file format used to read and write data tables | Apache Software Foundation | https://arrow.apache.org |
| DeepLabCut [version]: ResNet-50 pose-estimation network used to track the animal in the overhead video (model DLC_resnet50_cost_benefit_DLCFeb7shuffle1_900000) | (Mathis et al., 2018) | https://www.deeplabcut.org |
| Kilosort2 | (Pachitariu et al., 2016) | https://github.com/jamesjun/Kilosort2 |

## Supplemental table titles and legends

**Fig. S1 | Firing-rate dynamics, unit yield and area-pair coverage across the five recorded regions. a,** Heatmaps of trial-averaged, normalized single-unit firing rate for all recorded units in each of five regions (rows, top to bottom: MO/VO, ACC, DMS, VP, STA), shown across six aligned task epochs (columns, left to right): Pre-Cue, Offer, Action, Corridor Lg1, Corridor Lg6, and Pre-reward delay. Within each region, units are sorted by the time bin of peak normalized firing rate (a fixed sort order applied across all epoch columns). The vertical dashed line in each panel marks the alignment event for that epoch. Colour indicates normalized firing rate with lighter colours signifying higher normalised firing rate activity (rocket colormap; shared colorbar shown at the right of the DMS row). **b,** Distribution of per-session single-unit yield for each region, shown as a horizontal histogram (proportion of sessions per unit-count bin) with an overlaid Gaussian kernel density estimate and a dashed line marking the median yield. Panels are ordered and colour-coded as in panel a. **c**, Heatmap of the number of unique recording sessions per subject (columns) and area pair (rows) in which at least one unit passing the trial-averaged firing-rate criterion (>0.1 Hz) was recorded simultaneously in both regions of the pair, i.e. the sessions available for canonical correlation analysis of that pair. Cell values give the session count; colour (rocket colormap) reflects the same value. **d,** Combined unit yield per recording day for each pair of recorded areas (left) and pooled by circuit class (right). Each dot is one subject-day on which both areas of the pair were recorded, and its value is the number of units in the two areas combined, counting units that passed the firing-rate criterion. Boxes show the median and interquartile range, whiskers extend to 1.5 × the interquartile range, and box colour indicates circuit class (Fr-Fr, frontal–frontal; Fr-BG, frontal–basal ganglia; BG-BG, basal ganglia–basal ganglia).

**Fig. S2** | **Cross-animal reliability of the leading canonical correlation (CCA ρ₁) between area pairs. a,** Node-and-edge diagrams of the cross-validated leading canonical correlation (ρ₁) between each recorded area pair, shown separately for each animal (rows) across the six task epochs (columns). Nodes mark the five recorded regions at fixed anatomical schematic positions and are drawn (white, labelled by region) only where that region participates in at least one finite-valued edge in that panel. Edges connect area pairs for which a CCA model was successfully fitted; edge colour is a blend of the two connected regions’ assigned colours, edge width reflects the magnitude of ρ_1_ relative to other edges within that same animal (not directly comparable in absolute magnitude between rows), and each edge is annotated with its numeric ρ₁ value (Fisher z-averaged across sessions for that animal/epoch/area-pair). All finite-valued edges are shown regardless of statistical significance. **b,** Bootstrapped intra-class correlation coefficient (ICC) quantifying cross-animal agreement in ρ₁ for each area pair. For each pair, cross-validated ρ₁ values were pooled across all epochs and sessions and fit with a random-intercept mixed model (ρ₁ ∼ 1 + (1 | animal)); ICC = between-animal variance / within-animal variance, with lower ICC indicating greater cross-animal agreement. 95% confidence intervals were obtained by resampling sessions within animal (5,000 bootstrap resamples). For each area pair: black dot = point estimate, error bar = 95% bootstrap CI, coloured fill = full bootstrap ICC distribution (histogram bin width 0.025), dashed horizontal line = ICC of 1. Area pairs whose bootstrap distribution collapsed entirely to ICC = 1 (a degenerate mixed-model fit, arising when only one animal contributed data to that pair) are shown with their axis tick and animal count only, with no point, distribution, or error bar plotted. The number of animals (n) contributing to each area pair’s estimate is printed above the ICC = 1 reference line. **c,** Mean cross-validated leading canonical correlation (ρ₁) for each area pair: bar height represents the across-animal mean ρ₁ (averaged across epochs and sessions), with individual animals’ Fisher z-averaged ρ₁ overlaid as coloured scatter points, coloured to match panel a or b’s area-pair colour scheme.

**Fig. S3 | Coding axis comparisons, running-speed profiles, decoding in the corridor epochs, and accept–reject variability of the decoded task variables. a,** Axes of the variables that replace reward decoding are separate from earlier reward coding. For three area pairs (MO/VO-DMS, MO/VO-STA, ACC-DMS), points show the mean cosine between the decoding axis of the replacing task variable (velocity or trial count) in corridor epoch Lg1 or Lg6 and the reward decoding axis in each earlier epoch (colour: Offer, Action, Corridor Lg1). Axes were obtained by fitting one canonical correlation analysis per session, area pair and epoch on all valid trials, and mapping each variable’s decoder coefficients in canonical-component space back onto the units (u = W·b). Cosines were computed over the units present in both epochs’ analyses (≥5), separately for each area of the pair, and compared only where both variables were significantly decodable at the cohort level (p < .01). Points are means from a linear mixed model (random intercept per animal, session variance nested within animal) with 98% confidence intervals. The grey band marks the equivalence margin of ±0.2 (78.5°–101.5°). ≈, equivalent to zero (two one-sided tests, α = .01, Benjamini–Hochberg-corrected across all 24 comparisons); no cosine differed significantly from zero (Wald test, same correction). Unmarked intervals are inconclusive. b, Distribution of mean running velocity (cm/s) in each trial epoch, estimated from top-down DLC video tracking. Histograms are binned, and the x-axis cut off, at the 95th percentile of each epoch’s distribution; dashed lines mark the median. Insets (top right) show the average of the per-animal median velocity broken down by each trial’s reward and effort level (colour scale and values in cm/s, shared colorbar). c, Accepted-versus-rejected decoder comparison in the corridor epochs. Top: multiplex node diagrams for Corridor Lg1 and Lg6. An edge joins an area pair where the held-out decoding correlation (r) of a task variable differed between accepted and rejected trials (two-sided Wilcoxon signed-rank test of Δr against zero across sessions, p < .01, uncorrected). Edge colour indicates the decoded task variable, and area pairs significant for several variables carry parallel edges. Bottom: per-session Δr (accept − reject) pooled over the significant area pairs for each task variable. Bars show the mean and dots individual sessions. Task variables with no significant area pair in either epoch (effort) are omitted. d, Variability of the true (not decoded) task-variable values in accepted vs. rejected trials, For each session, the variance of each task variable’s true trial-level value was computed separately within accepted and rejected trials, after balancing the larger group down to the trial count of the smaller (mean of 20 random whole-trial subsamples). Bars show the across-session mean of log₂ (accept/reject variance), with positive values indicating more variable outcomes on accepted vs rejected trials. Asterisks mark task variables whose per-session ratios differ from zero (two-sided Wilcoxon signed-rank test, α = .01).

**Fig. S4 | The dominant CCA ρ is not significantly related to the trial-count coefficient of choice behaviour. a,** Fitted probability of accepting an offer against trial count for each session (thin lines, coloured by animal) and the cohort mean (thick black line). Curves come from per-session binomial GLMs of accept/reject choice on reward, effort and trial count, evaluated across each session’s observed trial range with reward and effort at the session median. The mean is taken up to the trial count reached by at least 75% of sessions. Sessions shown are those in **c,d** (n = 37). **b,** Slope of each session’s trial-count coefficient (change in log-odds of accepting per trial) on the session’s dominant CCA ρ, for each area pair. ρ_1_ is the held-out (10-fold cross-validated) first canonical correlation over the decision window, z-scored within area pair, so β is the change in the coefficient per 1 SD of ρ_1_. Slopes are from linear mixed models with a random intercept for animal. Bars show β ± S.E.M and are coloured by circuit class (Fr–Fr, BG–BG, Fr–BG). No slope reached p < .01 (uncorrected). **c,** ACC–STA ρ_1_ against the trial-count coefficient, the pair with the smallest p-value in **b** (37 sessions, 4 animals; β = −0.0015 per SD of ρ, 95% CI −0.0034 to +0.0004, p = .13, uncorrected). Dots are sessions coloured by animal; the line and shading are the population-level fit and 95% CI of the mixed model in **b**. **d,** ACC–STA dominant ρ across quartiles of the trial-count coefficient, with sessions pooled across animals. Boxes show the median and interquartile range, whiskers extend to 1.5 × the interquartile range, and dots are sessions coloured by animal.

**Table S1 | Edge values underlying the per-animal network diagrams in Fig. S2a.** Cross-validated canonical correlation (ρ_₁_; first canonical dimension, k ≤ 10) between each pair of recorded areas, for each animal (rows) and task epoch (sub-rows), as annotated on the corresponding edges of Fig. S2a. Values are Fisher z-averaged across that animal’s recording sessions and are not corrected for chance level (cf. Δρ in the main figure). Area pairs are organised along columns: MO/VO, motor/ventral orbital cortex; ACC, anterior cingulate cortex; DMS, dorsomedial striatum; VP, ventral pallidum; STA, subthalamic area. Dashes indicate area pairs for which an animal contributed no simultaneously recorded units.

