## Supplemental Information for "Dynamic information sharing across frontal-basal ganglia neurons during cost-benefit decision-making and reward pursuit"

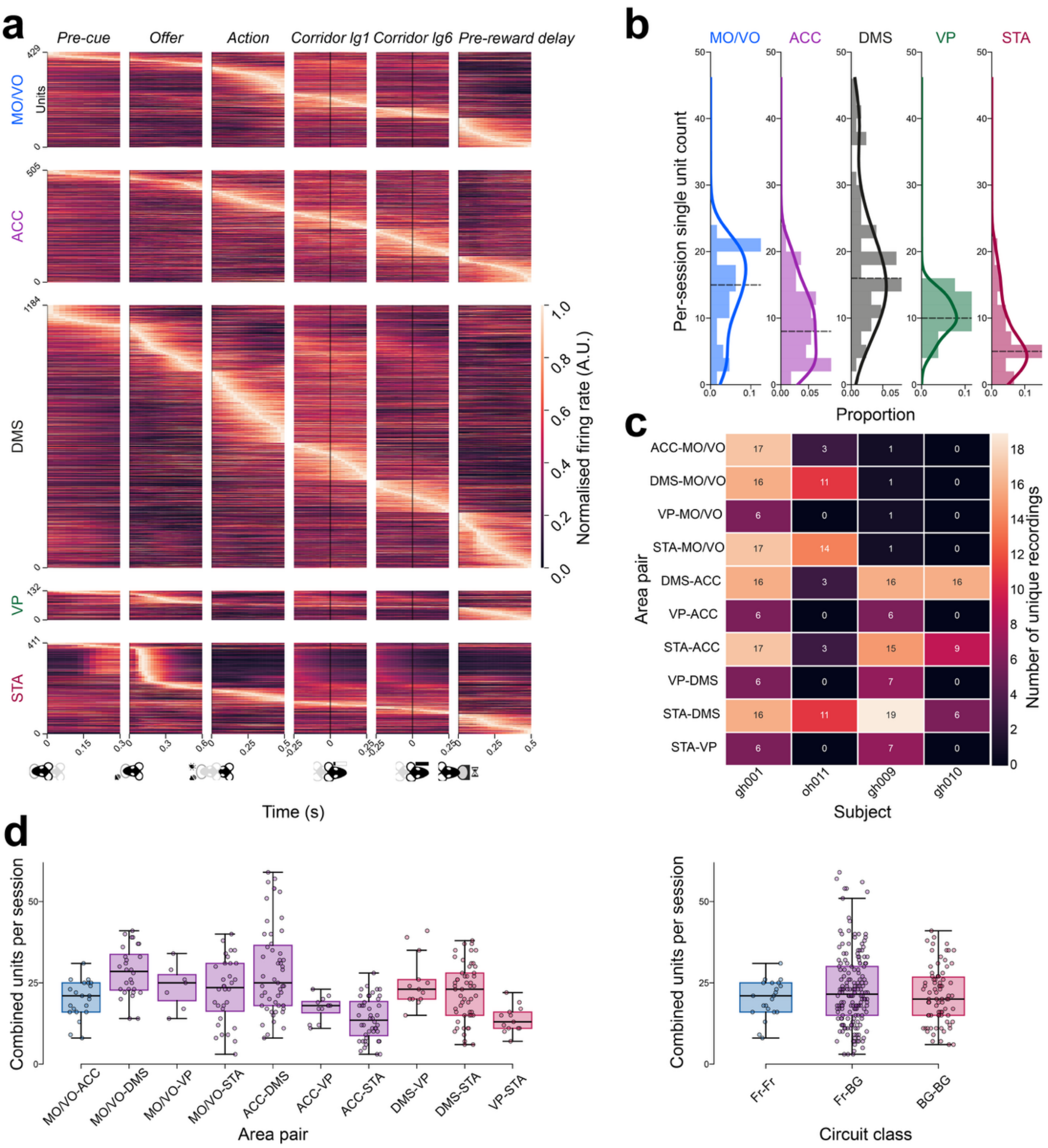


**Fig. S1 | Firing-rate dynamics, unit yield and area-pair coverage across the five recorded regions. a,** Heatmaps of trial-averaged, normalized single-unit firing rate for all recorded units in each of five regions (rows, top to bottom: MO/VO, ACC, DMS, VP, STA), shown across six aligned task epochs (columns, left to right): Pre-Cue, Offer, Action, Corridor Lg1, Corridor Lg6, and Pre-reward delay. Within each region, units are sorted by the time bin of peak normalized firing rate (a fixed sort order applied across all epoch columns). The vertical dashed line in each panel marks the alignment event for that epoch. Colour indicates normalized firing rate with lighter colours signifying higher normalised firing rate activity (rocket colormap; shared colorbar shown at the right of the DMS row). **b,** Distribution of per-session single-unit yield for each region, shown as a horizontal histogram (proportion of sessions per unit-count bin) with an overlaid Gaussian kernel density estimate and a dashed line marking the median yield. Panels are ordered and colour-coded as in panel a. **c**, Heatmap of the number of unique recording sessions per subject (columns) and area pair (rows) in which at least one unit passing the trial-averaged firing-rate criterion (>0.1 Hz) was recorded simultaneously in both regions of the pair, i.e. the sessions available for canonical correlation analysis of that pair. Cell values give the session count; colour (rocket colormap) reflects the same value. **d,** Combined unit yield per recording day for each pair of recorded areas (left) and pooled by circuit class (right). Each dot is one subject-day on which both areas of the pair were recorded, and its value is the number of units in the two areas combined, counting units that passed the firing-rate criterion. Boxes show the median and interquartile range, whiskers extend to 1.5 × the interquartile range, and box colour indicates circuit class (Fr-Fr, frontal–frontal; Fr-BG, frontal–basal ganglia; BG-BG, basal ganglia–basal ganglia).

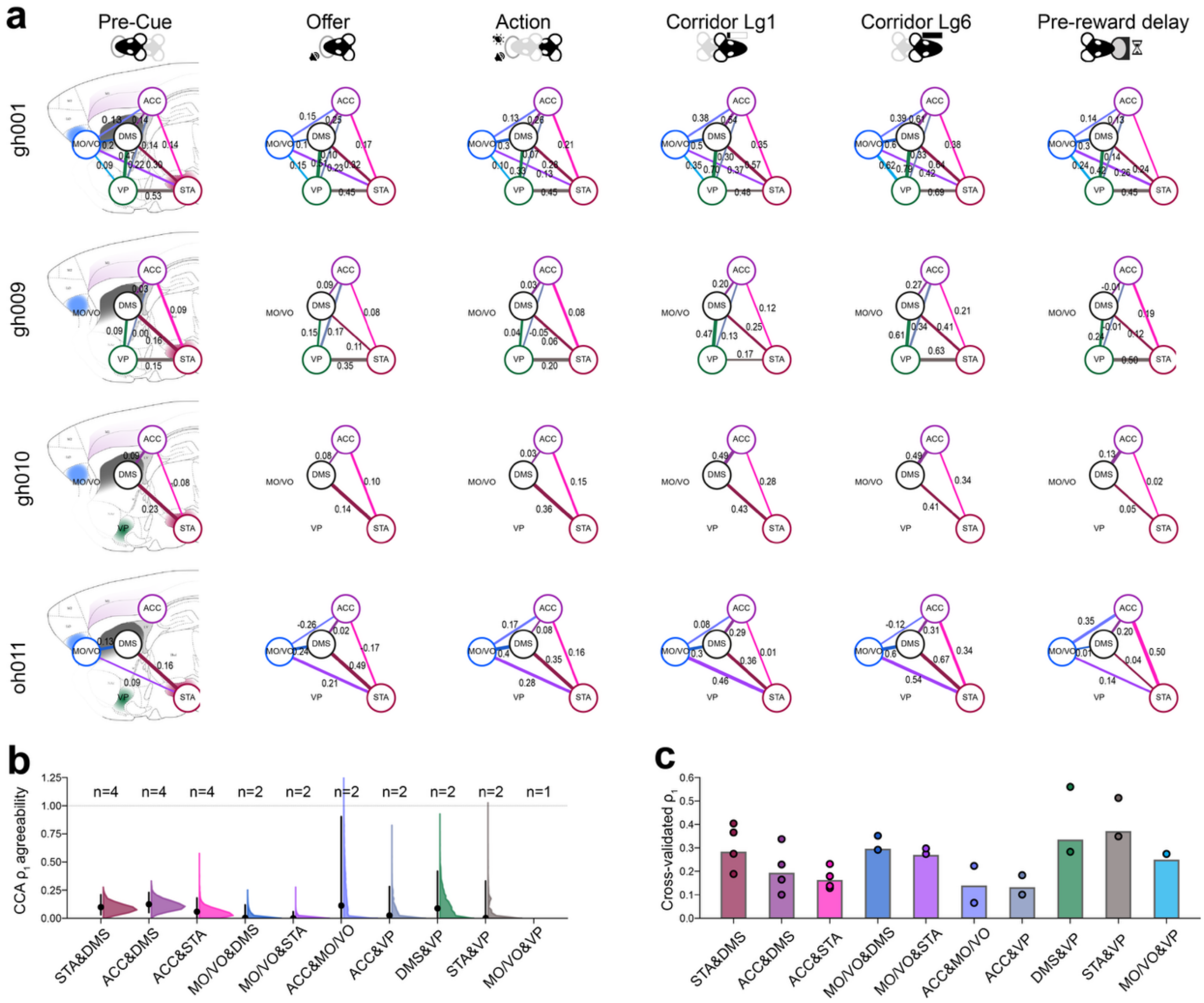


**Fig. S2 | Cross-animal reliability of the leading canonical correlation (CCA $\rho_1$) between area pairs.** **a,** Node-and-edge diagrams of the cross-validated leading canonical correlation ($\rho_1$) between each recorded area pair, shown separately for each animal (rows) across the six task epochs (columns). Nodes mark the five recorded regions at fixed anatomical schematic positions and are drawn (white, labelled by region) only where that region participates in at least one finite-valued edge in that panel. Edges connect area pairs for which a CCA model was successfully fitted; edge colour is a blend of the two connected regions' assigned colours, edge width reflects the magnitude of $\rho_1$ relative to other edges within that same animal (not directly comparable in absolute magnitude between rows), and each edge is annotated with its numeric $\rho_1$ value (Fisher z-averaged across sessions for that animal/epoch/area-pair). All finite-valued edges are shown regardless of statistical significance. **b,** Bootstrapped intra-class correlation coefficient (ICC) quantifying cross-animal agreement in $\rho_1$ for each area pair. For each pair, cross-validated $\rho_1$ values were pooled across all epochs and sessions and fit with a random-intercept mixed model ($\rho_1 \sim 1 + (1 \mid \text{animal})$); ICC = between-animal variance / within-animal variance, with lower ICC indicating greater cross-animal agreement. 95% confidence intervals were obtained by resampling sessions within animal (5,000 bootstrap resamples). For each area pair: black dot = point estimate, error bar = 95% bootstrap CI, coloured fill = full bootstrap ICC distribution (histogram bin width 0.025), dashed horizontal line = ICC of 1. Area pairs whose bootstrap distribution collapsed entirely to ICC = 1 (a degenerate mixed-model fit, arising when only one animal contributed data to that pair) are shown with their axis tick and animal count only, with no point, distribution, or error bar plotted. The number of animals (n) contributing to each area pair's estimate is printed above the ICC = 1 reference line. **c,** Mean cross-validated leading canonical correlation ($\rho_1$) for each area pair: bar height represents the across-animal mean $\rho_1$ (averaged across epochs and sessions), with individual animals' Fisher z-averaged $\rho_1$ overlaid as coloured scatter points, coloured to match panel a or b's area-pair colour scheme.

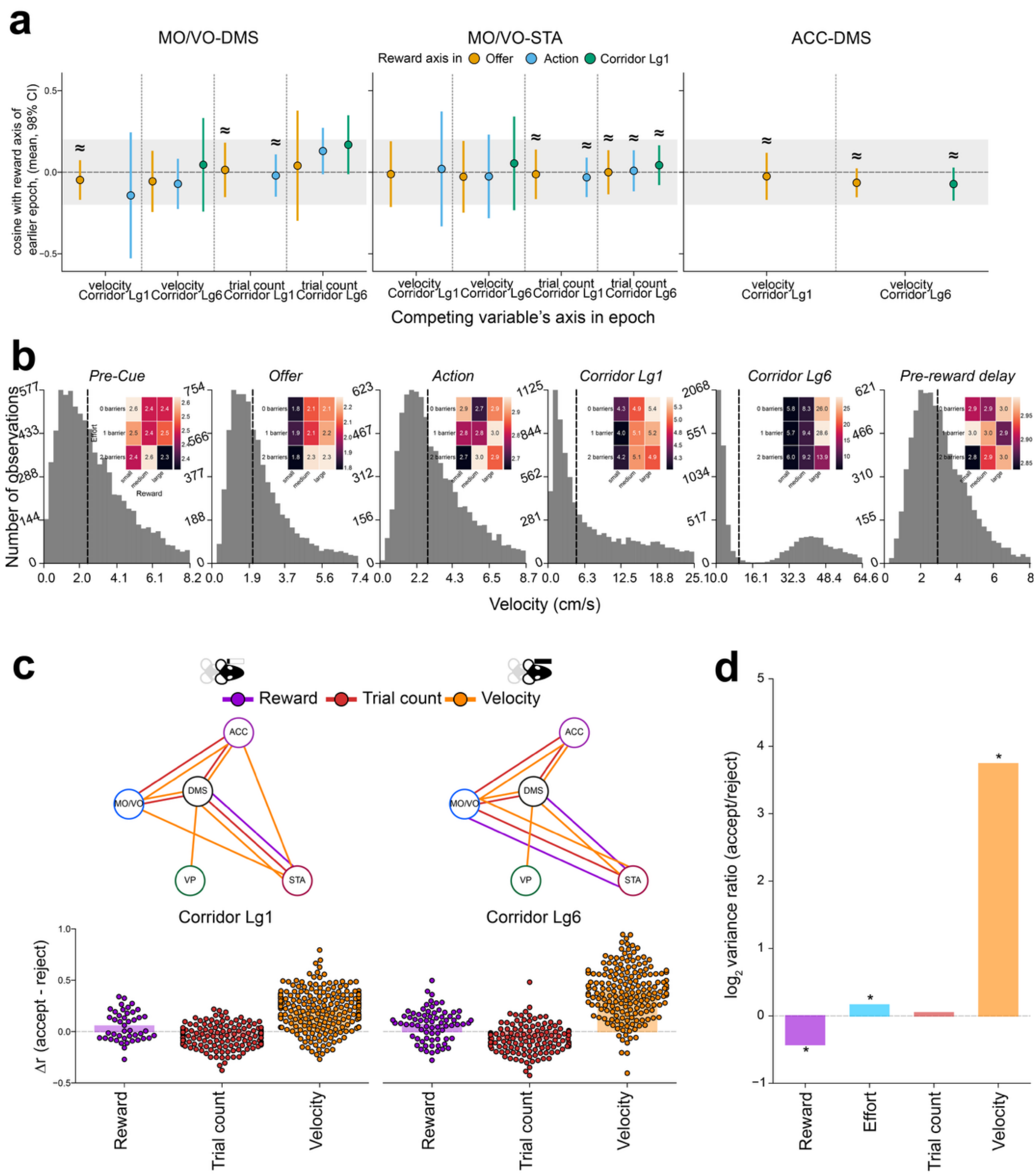


**Fig. S3 | Coding axis comparisons, running-speed profiles, decoding in the corridor epochs, and accept–reject variability of the decoded task variables. a,** Axes of the variables that replace reward decoding are separate from earlier reward coding. For three area pairs (MO/VO-DMS, MO/VO-STA, ACC-DMS), points show the mean cosine between the decoding axis of the replacing task variable (velocity or trial count) in corridor epoch Lg1 or Lg6 and the reward decoding axis in each earlier epoch (colour: Offer, Action, Corridor Lg1). Axes were obtained by fitting one canonical correlation analysis per session, area pair and epoch on all valid trials, and mapping each variable's decoder coefficients in canonical-component space back onto the units (u = W·b). Cosines were computed over the units present in both epochs' analyses (≥5), separately for each area of the pair, and compared only where both variables were significantly decodable at the cohort level ($p < .01$). Points are means from a linear mixed model (random intercept per animal, session variance nested within animal) with 98% confidence intervals. The grey band marks the equivalence margin of ±0.2 (78.5°–101.5°). ≈, equivalent to zero (two one-sided tests, $\alpha = .01$, Benjamini–Hochberg-corrected across all 24 comparisons); no cosine differed significantly from zero (Wald test, same correction). Unmarked intervals are inconclusive. **b,** Distribution of mean running velocity (cm/s) in each trial epoch, estimated from top-down DLC video tracking. Histograms are binned, and the x-axis cut off, at the 95th percentile of each epoch's distribution; dashed lines mark the median. Insets (top right) show the average of the per-animal median velocity broken down by each trial's reward and effort level (colour scale and values in cm/s, shared colorbar). **c,** Accepted-versus-rejected decoder comparison in the corridor epochs. Top: multiplex node diagrams for Corridor Lg1 and Lg6. An edge joins an area pair where the held-out decoding correlation (r) of a task variable differed between accepted and rejected trials (two-sided Wilcoxon signed-rank test of Δr against zero across sessions, $p < .01$, uncorrected). Edge colour indicates the decoded task variable, and area pairs significant for several variables carry parallel edges. Bottom: per-session Δr (accept − reject) pooled over the significant area pairs for each task variable. Bars show the mean and dots individual sessions. Task variables with no significant area pair in either epoch (effort) are omitted. **d,** Variability of the true (not decoded) task-variable values in accepted vs. rejected trials, For each session, the variance of each task variable's true trial-level value was computed separately within accepted and rejected trials, after balancing the larger group down to the trial count of the smaller (mean of 20 random whole-trial subsamples). Bars show the across-session mean of $\log_2$ (accept/reject variance), with positive values indicating more variable outcomes on accepted vs rejected trials. Asterisks mark task variables whose per-session ratios differ from zero (two-sided Wilcoxon signed-rank test, $\alpha = .01$).

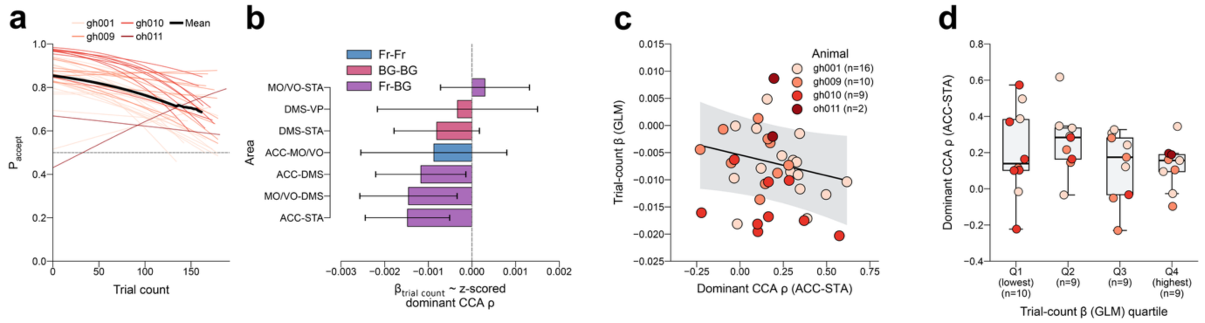


**Fig. S4 | The dominant CCA ρ is not significantly related to the trial-count coefficient of choice behaviour. a,** Fitted probability of accepting an offer against trial count for each session (thin lines, coloured by animal) and the cohort mean (thick black line). Curves come from per-session binomial GLMs of accept/reject choice on reward, effort and trial count, evaluated across each session's observed trial range with reward and effort at the session median. The mean is taken up to the trial count reached by at least 75% of sessions. Sessions shown are those in **c,d** ($n = 37$). **b,** Slope of each session's trial-count coefficient (change in log-odds of accepting per trial) on the session's dominant CCA ρ, for each area pair. $\rho_1$ is the held-out (10-fold cross-validated) first canonical correlation over the decision window, z-scored within area pair, so β is the change in the coefficient per 1 SD of $\rho_1$. Slopes are from linear mixed models with a random intercept for animal. Bars show β ± S.E.M and are coloured by circuit class (Fr–Fr, BG–BG, Fr–BG). No slope reached $p < .01$ (uncorrected). **c,** ACC–STA $\rho_1$ against the trial-count coefficient, the pair with the smallest p-value in **b** (37 sessions, 4 animals; $\beta = -0.0015$ per SD of ρ, 95% CI −0.0034 to +0.0004, $p = .13$, uncorrected). Dots are sessions coloured by animal; the line and shading are the population-level fit and 95% CI of the mixed model in **b**. **d,** ACC–STA dominant ρ across quartiles of the trial-count coefficient, with sessions pooled across animals. Boxes show the median and interquartile range, whiskers extend to 1.5 × the interquartile range, and dots are sessions coloured by animal.

**Table S1 | Edge values underlying the per-animal network diagrams in Fig. S2a.** Cross-validated canonical correlation ($\rho_1$; first canonical dimension, $k \leq 10$) between each pair of recorded areas, for each animal (rows) and task epoch (sub-rows), as annotated on the corresponding edges of Fig. S2a. Values are Fisher z-averaged across that animal's recording sessions and are not corrected for chance level (cf. $\Delta\rho$ in the main figure). Area pairs are organised along columns: MO/VO, motor/ventral orbital cortex; ACC, anterior cingulate cortex; DMS, dorsomedial striatum; VP, ventral pallidum; STA, subthalamic area. Dashes indicate area pairs for which an animal contributed no simultaneously recorded units.

| **Animal** | **Epoch** | **MO/VO–DMS** | **MO/VO–VP** | **MO/VO–STA** | **ACC–MO/VO** | **ACC–DMS** | **ACC–VP** | **ACC–STA** | **DMS–VP** | **STA–DMS** | **STA–VP** |
|---|---|---|---|---|---|---|---|---|---|---|---|
| **gh001** | Pre-cue | 0.18 | 0.09 | 0.22 | 0.13 | 0.14 | 0.14 | 0.14 | 0.47 | 0.30 | 0.53 |
| | Offer | 0.10 | 0.15 | 0.23 | 0.15 | 0.25 | 0.10 | 0.17 | 0.51 | 0.32 | 0.45 |
| | Action | 0.30 | 0.10 | 0.13 | 0.13 | 0.26 | 0.07 | 0.21 | 0.33 | 0.28 | 0.45 |
| | Corridor lg1 | 0.53 | 0.35 | 0.37 | 0.38 | 0.54 | 0.30 | 0.35 | 0.70 | 0.57 | 0.48 |
| | Corridor lg6 | 0.62 | 0.62 | 0.42 | 0.39 | 0.61 | 0.33 | 0.38 | 0.79 | 0.64 | 0.69 |
| | Pre-reward delay | 0.27 | 0.24 | 0.26 | 0.14 | 0.13 | 0.14 | 0.12 | 0.42 | 0.24 | 0.45 |
| **gh009** | Pre-cue | – | – | – | – | 0.03 | 0.00 | 0.09 | 0.09 | 0.16 | 0.15 |
| | Offer | – | – | – | – | 0.09 | 0.17 | 0.08 | 0.15 | 0.11 | 0.35 |
| | Action | – | – | – | – | 0.03 | -0.05 | 0.08 | 0.04 | 0.06 | 0.20 |
| | Corridor lg1 | – | – | – | – | 0.20 | 0.13 | 0.12 | 0.47 | 0.25 | 0.17 |
| | Corridor lg6 | – | – | – | – | 0.27 | 0.34 | 0.21 | 0.61 | 0.41 | 0.63 |
| | Pre-reward delay | – | – | – | – | -0.01 | -0.01 | 0.19 | 0.24 | 0.12 | 0.50 |
| **gh010** | Pre-cue | – | – | – | – | 0.09 | – | -0.08 | – | 0.23 | – |
| | Offer | – | – | – | – | 0.08 | – | 0.10 | – | 0.14 | – |
| | Action | – | – | – | – | 0.03 | – | 0.15 | – | 0.36 | – |
| | Corridor lg1 | – | – | – | – | 0.49 | – | 0.28 | – | 0.43 | – |
| | Corridor lg6 | – | – | – | – | 0.49 | – | 0.34 | – | 0.41 | – |
| | Pre-reward delay | – | – | – | – | 0.13 | – | 0.02 | – | 0.05 | – |
| **oh011** | Pre-cue | 0.13 | – | 0.09 | – | – | – | – | – | 0.16 | – |
| | Offer | 0.24 | – | 0.21 | -0.26 | 0.02 | – | -0.17 | – | 0.49 | – |
| | Action | 0.36 | – | 0.28 | 0.17 | 0.08 | – | 0.16 | – | 0.35 | – |
| | Corridor lg1 | 0.33 | – | 0.46 | 0.08 | 0.29 | – | 0.01 | – | 0.36 | – |
| | Corridor lg6 | 0.60 | – | 0.54 | -0.12 | 0.31 | – | 0.34 | – | 0.67 | – |
| | Pre-reward delay | 0.01 | – | 0.14 | 0.35 | 0.20 | – | 0.50 | – | 0.04 | – |
